# Developmental transcriptomics in *Pristionchus* reveals the logic of a plasticity gene regulatory network

**DOI:** 10.1101/2024.09.12.612712

**Authors:** Shelley Reich, Tobias Loschko, Julie Jung, Samantha Nestel, Ralf J. Sommer, Michael S. Werner

## Abstract

Developmental plasticity enables the production of alternative phenotypes in response to different environmental conditions. While significant advances in understanding the ecological and evolutionary implications of plasticity have been made, understanding its genetic basis has lagged. However, a decade of genetic screens in the model nematode *Pristionchus pacificus* has culminated in 30 genes which affect mouth-form plasticity. We also recently reported the critical window of environmental sensitivity, and therefore have clear expectations for when differential gene expression should matter. Here, we collated previous data into a gene-regulatory network (GRN), and performed developmental transcriptomics across different environmental conditions, genetic backgrounds, and mouth-form mutants to assess the regulatory logic of plasticity. We found that only two genes in the GRN (*eud-1* and *seud-1/sult-1*) are sensitive to the environment during the critical window. Interestingly, the time points of their sensitivity differ, suggesting that they act as sequential checkpoints. We also observed temporal constraint upon the transcriptional effects of mutating the GRN and revealed unexpected feedback between mouth-form genes. Surprisingly, expression of *seud-1/sult-1*, but not *eud-1*, correlated with mouth form biases across different strains and species. Finally, a comprehensive analysis of all samples identified metabolism as a shared pathway for regulating mouth-form plasticity. These data are presented in a Shiny app to facilitate gene-expression comparisons across development in up to 14 different conditions. Collectively, our results suggest that mouth-form plasticity evolved a constrained, two-tiered logic to integrate environmental information leading up to the final developmental decision.

## Introduction

Phenotypic plasticity is a widespread phenomenon by which exposure to different environments elicits different phenotypes from the same genotype (Pigliucci 2001; West-Eberhard 2003; DeWitt and Scheiner 2004). In sexually mature adults, plasticity is typically limited to physiological and behavioral traits, but when plasticity is channeled through development – often during an environmentally sensitive period or ‘window’ – dramatic differences in morphology can be achieved (Nijhout 2003). Several non-traditional model organisms which exhibit qualitative differences in morphology and behavior (i.e., polyphenism) have been developed to study plasticity, including social insects (Wilson 1971; Wheeler 1986; Simpson et al. 2011), dung beetles (Emlen 1994; Moczek 1998), and spadefoot toads (Pfennig 1990). These systems have revealed important roles for plasticity in evolution through its provision of environment-matched phenotypes and masking of genetic variants (West-Eberhard 1989; Pfennig et al. 2010; Moczek et al. 2011; Sommer 2020). However, to fully incorporate plasticity into the Modern Synthesis, a genetic framework is needed. Although possible (Kucharski et al. 2008; Kijimoto et al. 2012; Xu et al. 2015; Klein et al. 2016; Trible et al. 2017; Yan et al. 2017; Mazo-Vargas et al. 2017; Zhang et al. 2017; Parker and Brisson 2019), genetic analysis remains challenging in non-model organisms due to long reproductive cycles and difficulties in lab rearing.

The nematode *Pristionchus pacificus* was introduced as a model system for studying developmental plasticity in 2010 (Bento et al. 2010). *P. pacificus* has four larval stages (J1-J4) similar to *Caenorhabditis elegans*, although the first stage remains enclosed in the egg. The laboratory strain PS312 reaches sexual maturity within 72 h, at which point adults exhibit either a narrow, deep Stenostomatous (St) mouth form with a single dorsal tooth, or a wide, shallow Eurystomatous (Eu) mouth form containing two hooked teeth (**Fig. 1A & C**). This morphological decision also has ecological consequences as the St morph is limited to a bacterial diet, whereas the Eu morph is a facultative predator on other nematodes. Mouth-form development is sensitive to many environmental factors including pheromones, crowding, salt concentration, temperature, and culturing substrate (Bento et al. 2010; Bose et al. 2012; Ragsdale et al. 2013; Werner et al. 2017, 2018; Lenuzzi et al. 2021). Reciprocal transplantation experiments recently identified 36-60 h (J3-J4) as the critical window for mouth-form plasticity (Werner et al. 2023).

**Figure 1.**
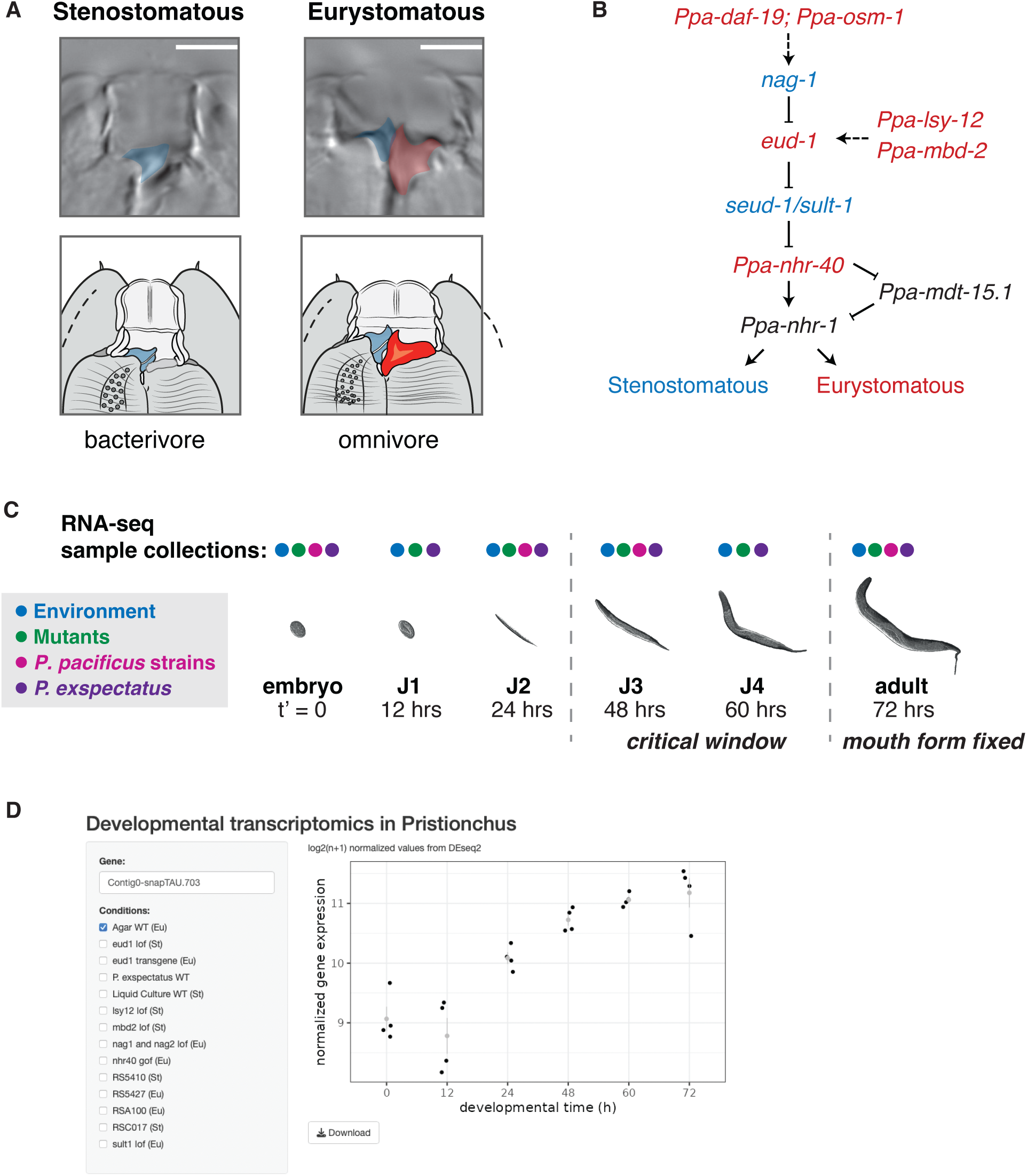
Developmental transcriptomics of mouth-form plasticity in *Pristionchus pacificus*. A) The two mouth-forms developed by adult *P. pacificus* with teeth colored for ease of visualization. B) A gene regulatory network for mouth-form development with St-promoting genes colored in blue and Eu-promoting genes in red. Genes in black may be downstream in both MF pathways. C) The life cycle of *P. pacificus* with colored circles showing the respective developmental time- points sampled for each condition. D) A screenshot of the Shiny app that can be used to visualize transcriptomic data for any gene in any conditions.

Additionally, more than a decade of sampling for *Pristionchus* has revealed a world-wide distribution with at least 48 species and hundreds of natural isolates (McGaughran et al. 2016; Kanzaki et al. 2021). Importantly, different natural isolates and species vary in the percentage of animals that are Eu or St, even when grown under the same environmental conditions (Ragsdale et al. 2013; Werner et al. 2017), demonstrating a genetic basis – and possible selection – for mouth-form ratios in the wild.

*P. pacificus* is easy to maintain in the lab and possesses many of the same traits that have made *C. elegans* a powerful model organism: hermaphroditism, short generation time, small genome, and amenability to genetic analysis. Over the last fourteen years, forward genetic screens and targeted mutagenesis in *P. pacificus* have identified many genes that affect mouth-from development (Bento et al. 2010; Ragsdale et al. 2013; Serobyan et al. 2016; Kieninger et al. 2016; Sieriebriennikov et al. 2017, 2018, 2020; Moreno et al. 2018, 2019; Namdeo et al. 2018; Bui et al. 2018; Lenuzzi et al. 2021; Sun et al. 2022a, 2022b; Casasa et al. 2023; Ishita et al. 2023; Levis and Ragsdale 2023). While there are undoubtedly more components involved, these experiments represent an important milestone as all phases of mouth-form development are now represented from environmental sensation to morphological execution of the decision. Furthermore, the identification of these genes enables mechanistic investigations into how plasticity works.

Previous studies have examined transcriptional differences between a few mouth-form mutants from mixed-staged populations. These data helped to identify downstream targets of mouth-form genes (Bui and Ragsdale 2019; Sieriebriennikov et al. 2020) and gene co-expression modules (Casasa et al. 2021). Here, we expand upon these studies by assembling a gene regulatory network (GRN) of mouth-form plasticity, and use developmental transcriptomics in different *Pristionchus* strains, species, and environmental conditions to identify 1) which genes in the network are environmentally responsive, 2) when they are responsive during development, and 3) how these patterns change over evolutionary time.

## Results

### A gene regulatory network for mouth-form plasticity

First, we performed a comprehensive literature analysis to compile all genes that have been shown to affect mouth form in *P. pacificus*. We identified 30 genes across 18 publications from 2010 to 2023. In **Table 1** we provide a list of these genes with their identifiers, chromosomal position, homolog in *C. elegans*, gene ontology, mouth-form phenotype and relevant citations. For the rest of this manuscript, we chose to focus on genes that have a ≥90% penetrant mouth-form phenotype when mutated. We ordered these genes into a gene regulatory network (GRN) for mouth-form plasticity using annotated protein function and previous epistatic experiments (**Fig. 1B**). At the start of the pathway are the chemosensory genes *daf-19* and *osm-1*, which control cilia development and function. The core of the network consists of a multi-gene locus containing the sulfatase *eud-1* and an antagonistic gene *nag-1*, two chromatin modifiers (*lsy-12*, *mbd-2*), and the sulfotransferase *seud-1/sult-1*. Downstream are two nuclear hormone receptors (*nhr-40, nhr-1*) and a Mediator subunit (*mdt-15.1*). This pathway, based on over a decade of forward and reverse genetic experiments, represents a comprehensive gene regulatory network of developmental plasticity.

**Table 1.**
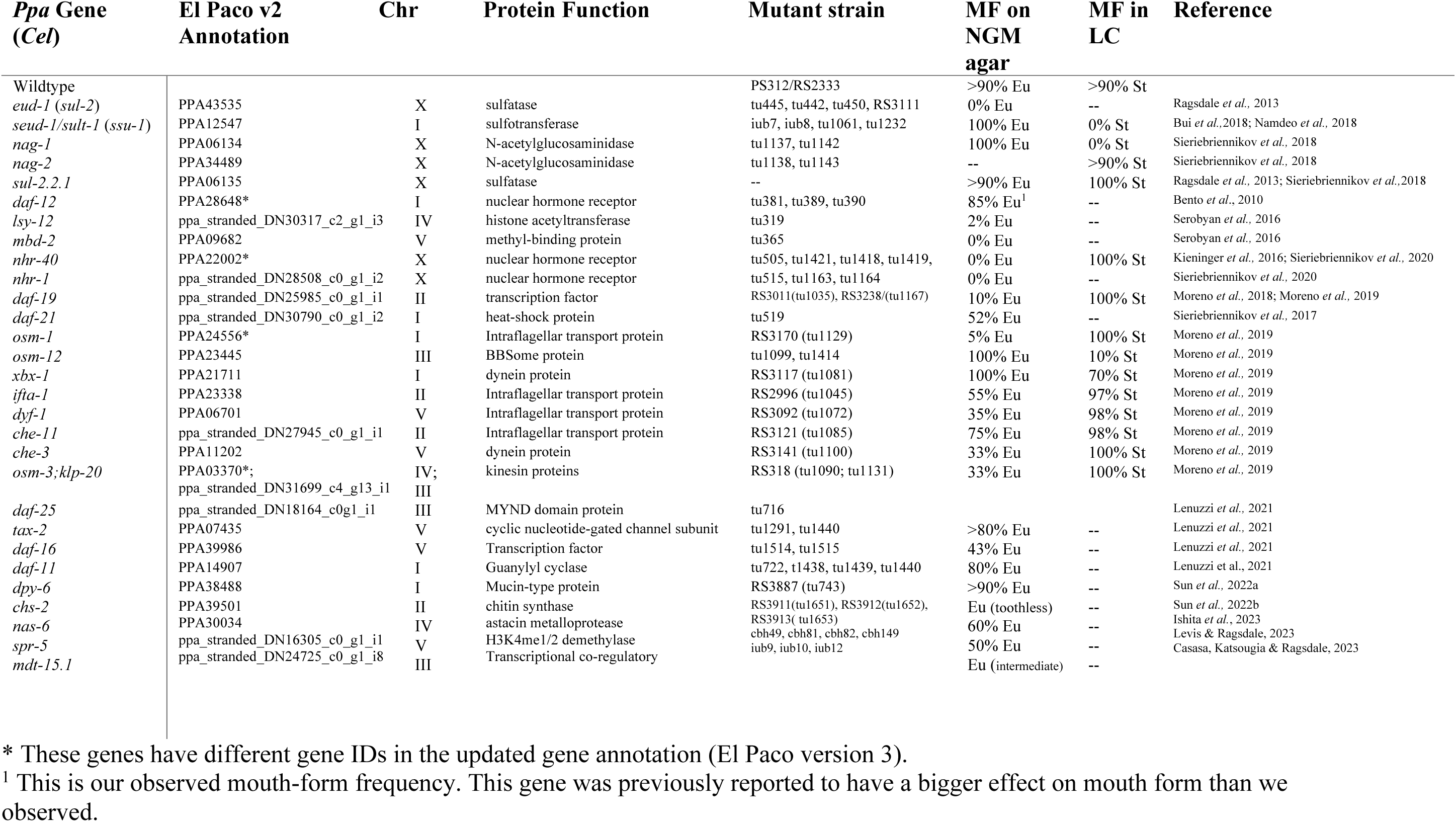
Genes shown to affect mouth-form development in *Pristionchus pacificus*. Gene IDs are based on the El Paco v2 gene annotation that was used for this transcriptomic analysis.

### Environmental responsiveness of the gene regulatory network for mouth-form plasticity

Developmental plasticity is mediated by environmental exposure during a sensitive period of development. Establishing the temporal action of molecular factors relative to this critical window is required to interpret the regulatory logic of developmental plasticity. We also sought to investigate potential feedback loops, and to understand how this logic evolves. To address these questions, we generated a developmental transcriptomics dataset that incorporates two environmental conditions, seven mouth-form mutant strains, four natural isolates of *P. pacificus*, and its sister species, *P. exspectatus* (**Supplemental Fig. 1A**). For each condition, we collected worms at multiple timepoints across development from 0 h post egg-synchronization (embryos) to 72 h (young adults) (**Fig. 1C**). We extracted and sequenced mRNA to a minimum depth of 3 million reads and with a minimum quality score of 30, ultimately resulting in over 3.5 billion high quality reads that we anticipate will be a valuable resource for the *P. pacificus* and phenotypic plasticity communities (**Supplemental Fig. 1-7, Supplemental Tables 1 & 4**). To facilitate visualization of these data, we made a Shiny app in R which renders downloadable plots of gene expression for any *P.pacificus* gene in each of the tested conditions (**Fig. 1D**, Chang et al. 2024). This app can be found at: https://pristionchus-transcriptomics.shinyapps.io/GRN-shiny/.

First, we analyzed how the GRN is affected by the environment. We cultured the *P. pacificus* laboratory strain PS312 on NGM-agar plates, which yield ∼95% Eu animals, and in S-Medium liquid culture, which yields 5-10% Eu animals (Werner et al. 2017), and collected samples at six developmental timepoints (0h, 12h, 24h, 48h, 60h, and 72 h, **Supplemental Fig. 8**). We then performed polyA-tail based bulk RNA-seq and used DESeq2 (Love et al. 2014) to assess differential gene expression at each time point. A principal component analysis of normalized expression showed that PC1, which separated samples by developmental time, explained 88% of the variance in gene expression (**Supplemental Fig. 9A)**. This suggests that developmental time is the single largest factor that explains differences in gene expression and highlights the importance of using stage-synchronized samples for transcriptomic analysis.

Overall, we found a total of 7,648 differentially expressed genes (DEGs) between agar and liquid culture samples across all time points. In general, the number of DEGs between the two conditions increased with developmental time. The 12 h timepoint, which contains a mix of eggs and J1s, had only 81 DEGs (1% of all DEGs) (**Fig. 2A**). At 24 and 48 h there was a notable increase to 1,255 and 842 DEGs, respectively (**Fig. 2B & C**). Yet, at 60 and 72 h time points there were far more DEGs (4937 and 4640, respectively, **Fig. 2D & E**, **Supplemental Fig. 9B & 9C**). Slightly fewer than half of the DEGs genes were differentially expressed at multiple developmental timepoints, and the last two developmental timepoints shared >60% (2,995) of their DEGs (**Supplemental Fig. 9C**). Altogether, these data show a dramatic transcriptomic response to the different environmental conditions starting between 48-60 h that persists through to adulthood. Interestingly, this transcriptomic response coincides with the closing of the critical window for mouth-form plasticity (Werner et al. 2023). We conclude that the early stages of development are relatively canalized with only minor changes in gene expression occurring in response to the environment. Leading up to the end of the critical window, large-scale transcriptional rewiring occurs in response to the environment, which bifurcates development into different trajectories, including mouth- form development. Presumably, the mouth-form genes that are differentially expressed early in the critical window represent environmental switch genes, which if expressed above a threshold initiate the large-scale transcriptional changes observed at the end of the critical window (*i.e.,* ≥60 h).

**Figure 2.**
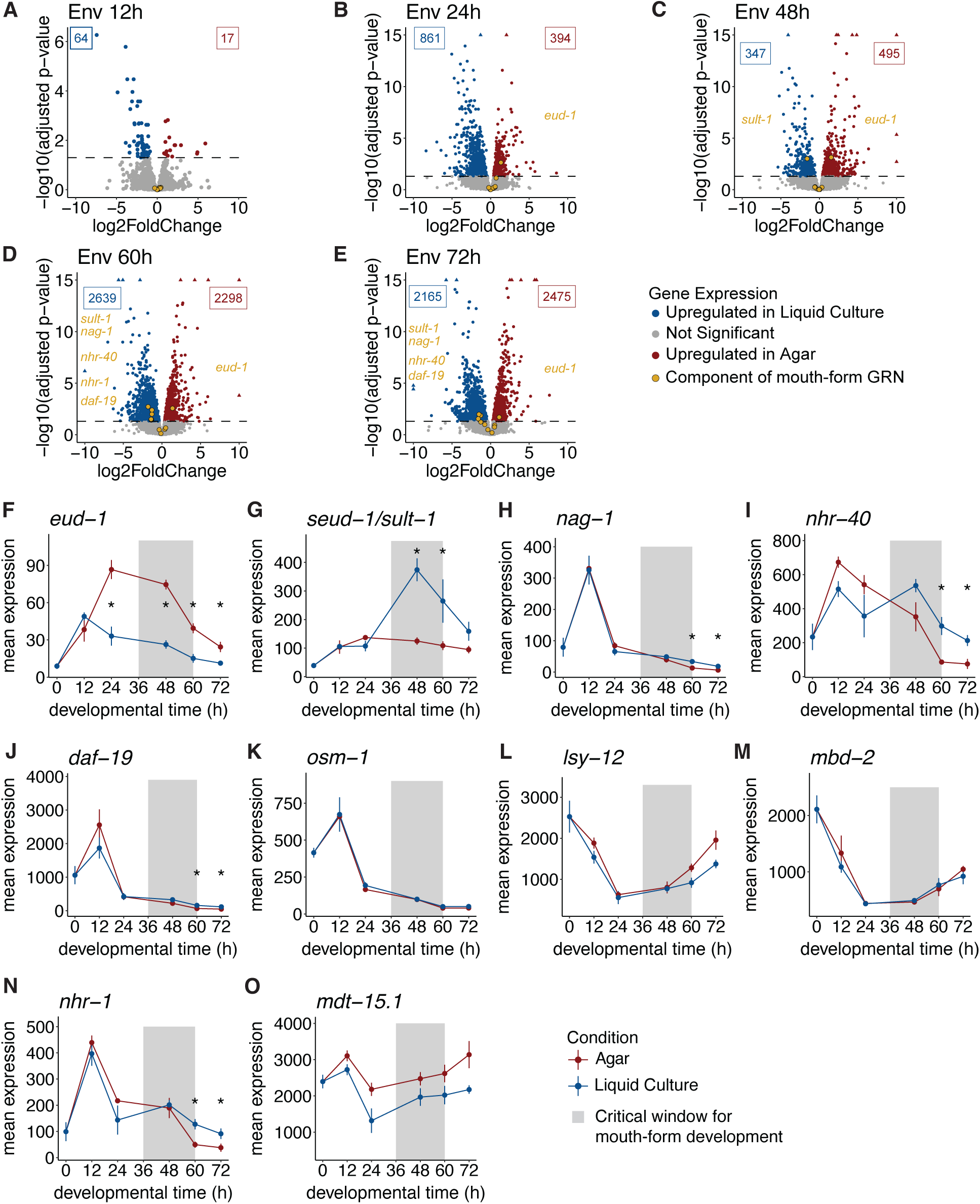
A subset of the mouth-form GRN is environmentally responsive. A-E) Volcano plots of genes differentially expressed between worms grown on agar and in liquid culture at five developmental time- points. Significant genes (adjusted p-values <0.05) are colored by the condition in which they were up- regulated, blue for liquid culture and red for agar. Siginificant mouth-form genes are labelled on the plot in gold. F-O) Mean expression of normalized counts from DESeq2 with SEM for 10 mouth-form genes from the GRN. * designates time points with significant differences in expression (adjusted p-value <0.05).

Next, we investigated whether the environment induces changes in expression of the mouth-form GRN, and whether any members are differentially expressed before the critical window. We were surprised to find that even though each of these genes have a nearly 100% penetrant phenotype when mutated, only six of the ten genes were significantly differentially expressed (*adjusted p-value<*0.05) at any point during development (**Fig. 2F-O**). Four of these genes exhibited a change in expression that is consistent with their loss-of-function phenotype. Curiously though, *nhr-40* and *daf-*19, which are thought to promote the Eu mouth form based on their loss-of-function phenotypes (**Table 1**, Moreno *et al*. 2019; Sieriebriennikov *et al*. 2020), were upregulated in St-promoting liquid culture during late development (**Fig. 2I & J**). While the basis of the discrepancy is not currently clear, it is possible that the nuclear hormone receptor *nhr-40* may be downstream of both mouth-form pathways (see results from mutant comparisons in following section).

All the differentially expressed mouth-form genes showed differences in expression at the 60 h time point, and four of these genes continued to show differences in expression into adulthood. At 60 h animals are almost exclusively juvenile stage 4 (J4) in both conditions, which means they still go through one more molt before adulthood. Hence it is possible that these differences contribute to the mouth-form decision. However, as the critical window has largely closed by 60 h, we suspect that these genes do not contribute to the switch mechanism *per se*. Instead, either their function later in development is unrelated to their role in mouth form, or late differential expression pertains to their role in executing the mouth-form decision. For example, *nhr-1* exhibits late differential expression (**Fig. 2N**) and yields intermediate mouth forms when mutated (Sieriebriennikov et al. 2020); results consistent with it regulating the expression of additional genes required for the formation of the adult mouth.

In contrast, the genes *eud-1* and *seud-1/sult-1* showed significant differences in expression as early as 24 and 48 h, respectively (**Fig. 2F & 2G**). These data support the role of *eud-1* and *seud-1/sult-1* as opposing genetic switches for mouth-form development, and the downstream placement of the nuclear hormone receptors. Moreover, the early differential expression of *eud-1* at 24 h – the only mouth-form gene to be differentially expressed prior to the critical window – suggests that it is the pivotal environmental switch gene.

### Transcriptional effects of mutations in the mouth-form GRN are temporally restricted

To explore the relationship between these mouth-form genes, we performed RNA-seq on seven mutant lines of *P. pacificus* grown on NGM agar plates (**Supplemental Fig. 1A**) and quantified changes in gene expression relative to wildtype PS312. We reasoned that keeping the environment consistent across samples should yield fewer changes in gene expression than between-environment comparisons.

However, we found that mutating or over-expressing single mouth-form genes generated substantial transcriptomic differences across development, especially at 60 and 72 h timepoints (**Supplemental Fig. 10A**). Nonetheless, when we pooled all mutant strains together and compared between Eu and St samples, we found a reduction in the number of differentially expressed genes across development (**Supplemental Fig. 10A**). This indicates that while each individual mouth-form mutant exerts large transcriptional changes compared to wildtype, there are relatively few genes that are differentially regulated in common.

As mouth-form dimorphism represents an evolutionary novelty in Diplogastrids, the large differences in gene expression seen by individual mutants suggest that the mouth-form GRN still has ancestral roles not involved in mouth form. This interpretation is consistent with the hypothesis that plasticity evolves by co-opting genes from pre-existing networks, as previously suggested by the enrichment of starvation and dauer-related genes among mouth-form specific gene co-expression modules (Casasa et al. 2021).

Notably, the *eud-1* loss-of function mutant had the fewest DEGs and appears to be an outlier in this regard (Z= −1.9, **Supplemental Fig. 10B**). Collectively, a meta-analysis of GRN mutants across development suggests that *eud-1* is the most specific component of mouth-form regulation.

We next investigated the transcriptional effects of mutating individual mouth-form genes on other components of the GRN. First, we found that mutations in mouth-form genes supported the overall topology of the GRN (**Fig. 1B**). For example, overexpression of *eud-1* led to upregulation of the two downstream nuclear hormone receptors (**Fig. 3B & 3C**), whereas a *loss-of-function* mutation in *eud-1* resulted in an upregulation of *seud-1/sult-1* (**Fig. 3D**). Conversely, mutating *seud-1/sult-1* had no effect on *eud-1* expression (**Fig. 3G**), supporting its downstream placement in the GRN relative to *eud-1*. The feedback from *eud-1* to *seud-1/sult-1* was notable because neither of their protein products are transcription factors, and they are believed to be expressed in different cells (Bui et al. 2018). Thus, there must be indirect crosstalk between switch genes.

**Figure 3.**
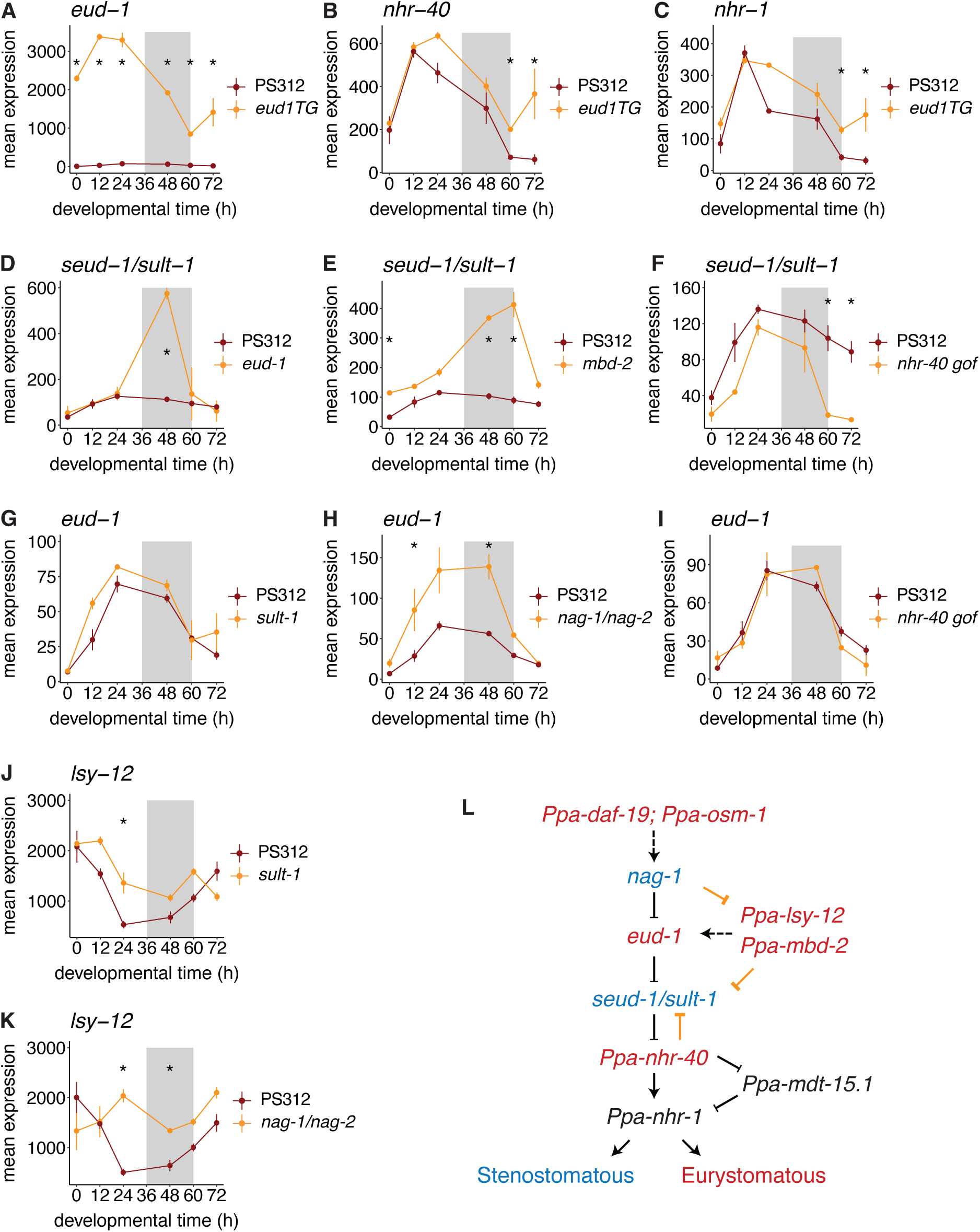
Transcriptional effects of mutations in the mouth-form GRN are temporally restricted. A-K) Mean expression from DESeq2 normalized counts of mouth-form genes in wildtype PS312 and indicated mutant line. * indicates time point with significant differences in expression (adjusted p-value <0.05). L) Mouth form GRN with new regulatory links drawn in gold.

*eud-1* is part of a multi-gene locus and is bookended by two genes *nag-1* and *nag-2*, which promote the St morph. The *nag-1/2* double mutant showed an upregulation of *eud-1* across development with significant differences from wildtype at 12 h and 48h (**Fig. 3H**), supporting its placement upstream of *eud-1*. Similar to *eud-1* and *seud-1/sult-1*, *nag-1/2* are cytosolic enzymes (N-acetylglucosaminidases). Thus, *nag-1/2, eud-1* and *seud-1/sult-1* exhibit regulatory connections with each other - presumably mediated via their enzymatic products functioning as signaling molecules.

Second, we found evidence for additional regulation between components of the mouth-form GRN. The *mbd-2* mutant displayed dramatic upregulation of *seud-1/sult-1* at 48 and 60 h (**Fig. 3E**). *mbd-2* is a methyl-binding protein that regulates *eud-1*’s expression (Serobyan et al. 2016) and may thus affect *seud- 1/sult-1*’s expression indirectly via *eud-1* or may directly regulate *seud-1/sult-1*’s expression. We also noted upregulation of *lsy-12* at 12 h in the *sult-1* mutant (**Fig. 3J**) and at 24 and 48 h in the *nag-1/2* double mutant (**Fig. 3K**). Finally, our dataset indicated a potential feedback mechanism from a downstream nuclear hormone receptor to an upstream switch gene. The gain-of-function mutation in *nhr- 40* caused a dramatic downregulation of *seud-1/sult-1*, but not *eud-1*, at 60 and 72 h compared to wildtype (**Fig. 3F & 3I**). These results enabled us to add new regulatory connections to the mouth-form GRN (**Fig. 3L**).

Third, we were struck by the temporal restriction of mouth-form mutants on other mouth-form genes. For example, even though *eud-1* was dramatically overexpressed at each sampled timepoint in the transgenic line (**Fig. 3A**), its upregulation of the downstream nuclear hormone receptors was limited to the last two timepoints (60 h and 72 h, **Fig. 3B & 3C**). Similarly, the upregulation of *seud-1/sult-1* in the *eud-1* mutant was limited to the 48 h timepoint (**Fig. 3D**), and the gain-of-function mutation in *nhr-40* resulted in upregulation of both nuclear hormone receptors, but only at 60 and 72 h (**Supplemental Fig. 11A & G**). Curiously, the nuclear hormone receptor *nhr-40* was upregulated in mutant samples relative to wildtype regardless of whether the mutant was Eu or St, perhaps suggesting that this gene may be involved in the determination of both mouth-forms (**Supplemental Fig. 11B-F**). In support of this hypothesis, *nhr-40* loss-of-function mutants are St, whereas gain-of-function mutants are Eu (Kieninger et al. 2016; Sieriebriennikov et al. 2020). Thus, our transcriptomic analyses indicate dynamic feedback across development between genes in a plasticity network. Surprisingly, this regulation is largely confined to the time points at which they are environmentally responsive, regardless of whether they function in the switch mechanism or the execution of the decision. This constraint is consistent with transcriptional regulation being confined to discrete developmental checkpoints and the critical window.

### Genetic background has different effects on the GRN than the environment

Natural isolates of *Pristionchus pacificus* vary in the ratio of Eu to St animals even when grown under the same environmental conditions, indicating a strong effect of genetic background on mouth-form development (Ragsdale et al. 2013). A recent study employed recombinant inbred lines and QTL mapping with two strains from clade B (Dardiry et al. 2023), and found copy-number variation of *cis*-regulatory elements underlying differences in *eud-1* expression and mouth form. However, outside of this case study, it is unknown which nodes of the network are being acted on by evolution. To expand upon these findings, we compared gene expression between two Eu- (RSA100 and RS5427) and two St-biased (RSC017 and RS5410) strains of *P. pacificus* from clade C. These strains were all grown on NGM-agar plates, and we isolated mRNA from samples at 0 h, 24 h, 48 h, and 72 h.

We observed that both St strains exhibited elevated *seud-1/sult-1* expression relative to St strains, whereas we did not observe the reciprocal pattern for *eud-1* (**Supplemental Fig. 12**). To increase our statistical power, we pooled the two Eu-biased strains and two St-biased strains. Overall, we found relatively few DEGs compared to the number of genes that were differentially expressed between environmentally-induced Eu and St worms (**Fig. 4A-D**). The greatest number of differentially expressed genes between Eu and St-biased strains was at the 0 h time point (**Fig. 4A**). PCA between natural isolates suggested that these differences are not due to differences in developmental rate (**Supplemental Fig. 13**), and a gene ontology revealed a broad distribution of cellular functions (**Supplemental Table 2**). These results suggest that Eu and St-biased strains are primed for differential developmental trajectories from the earliest stages of development. Consistent with the environmental comparisons, the 72 h time point exhibited substantially more DEGs than 48 h or 24 h (>2.5 times as many DEGs as the 48 h time point) (**Fig. 4C & D**), suggesting that distinct transcriptional changes accompany the formation of the two different adult mouth forms. Intriguingly, in this comparison only two of the genes from the mouth-form GRN showed significant differences in expression: *seud-1/sult-1* and *nag-1* (**Fig. 4F & G**), while *eud-1* did not show significant differences in expression at any time point (**Fig. 4E**). Thus, the environment and genetic background affect the mouth-form GRN differently. The significant difference in *seud-1/sult-1*’s expression was at 48 h, essentially in the middle of the critical window, whereas *nag-1* was differentially expressed at 72 h. This temporal pattern, and the absence of other differentially expressed mouth-form genes, suggests that *seud-1/sult-1* is the key GRN component responsible for the differences in mouth- form between the four natural isolates in this analysis.

**Figure 4.**
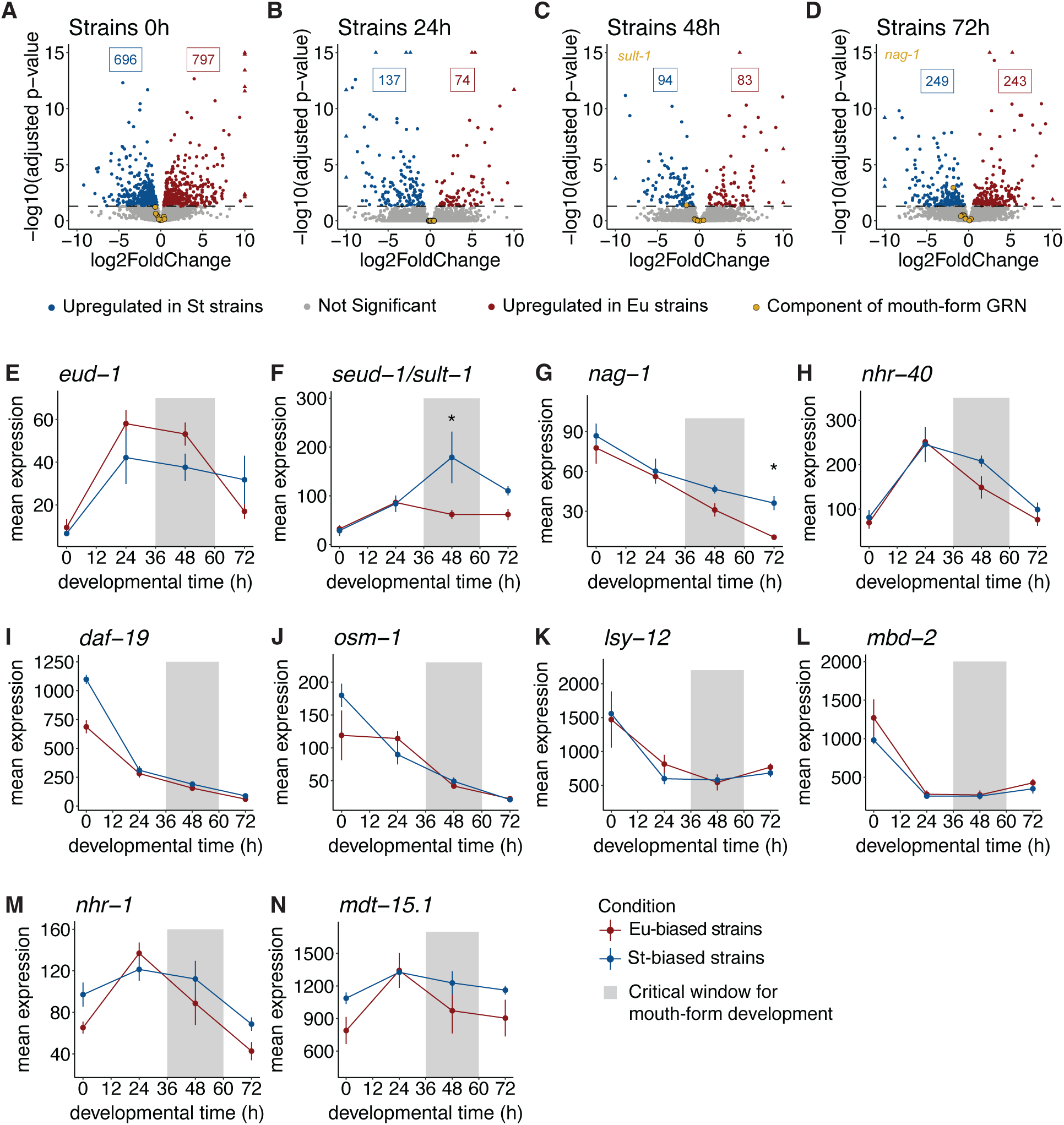
Natural variation in mouth-form transcriptomics. A-D) Volcano plots of differentially expressed genes between pooled St- and Eu-biased strains of *P. pacificus* at four developmen- tal timepoints. Significant genes (adjusted p-values <0.05) upregulated in St are colored blue and those upregulated in Eu are colored red. E-N) Mean expression of normalized counts from DESeq2 for mouth-form genes in pooled St- and Eu-biased strains. * designates time points with significant expression differences (adjusted p-value <0.05).

### *Pristionchus* sister species exhibit large transcriptomic differences

To further investigate the effect of genetic background on the mouth-form GRN, we compared gene expression between two sister species of *Pristionchus*, both of which exhibit mouth-form plasticity but with opposite biases in mouth-form development. In contrast to the main laboratory strain of *P. pacificus* (PS312), which develops primarily as Eurystomatous on agar plates (>95% Eu), the lab strain of *P. exspectatus* (RS5522B) develops almost completely as Stenostomatous (<1% Eu) (Kanzaki et al. 2012; Ragsdale et al. 2013). Evaluating gene expression between species is notoriously challenging (Munro et al. 2022). We took a conservative approach and mapped the reads of each species to its own transcriptome and then performed a differential expression analysis exclusively on the 1:1 species orthologs using DESeq2 (see Methods). We found that *P. pacificus* and *P. exspectatus* have large differences in gene expression across all developmental time points (**Supplementary** Fig. 14A-F).

Principal component analysis of the normalized expression values separated samples first by developmental time along PC1 and second by species along PC2 (**Supplemental Fig. 15**). The *mbd-2* gene in *P. exspectatus* seems to be incorrectly annotated and was excluded from this analysis. Every other mouth-form gene, with the exception of *nhr-1* (adjusted p-value = 0.056), was differentially expressed at some point during development (**Supplementary** Fig. 14G-O). Surprisingly, only two mouth-form genes, *seud-1/sult-1* and *nag-1*, showed significant differences during the critical window for the mouth-form decision, and in the expected direction (i.e., upregulated in *P. exspectatus* at 48 h) (**Supplementary** Fig. 14H **& I**). *eud-1* was differentially expressed between the two species, but only at 12 h and was unexpectedly upregulated in the St-biased *P. exspectatus* (**Supplementary** Fig. 14G). This result is in contrast to prior work showing *eud-1* to be upregulated in *P. pacificus* relative to *P. exspectatus* (Ragsdale et al. 2013). This could be due to differences in the annotations or pipelines used for the different experiments. Ragsdale *et al*. used FPKM to quantify gene expression relative to all genes and did not test for significance between the two species. Interestingly, two paralogs of *eud-1* were also significantly upregulated in *P. exspectatus* at multiple timepoints across development (**Supplemental Fig. 14P & Q**). Thus, comparing gene expression between sister species shows a massive rewiring of transcription – yet regulation of specific components of the plasticity GRN appear to be maintained to effect mouth-form biases.

Taken together, a comparison of mouth-form induction by the environment versus two scales of genetic background (strain and species) indicates some similarities and some differences. While many components of the GRN are differentially expressed between species, increasing or decreasing *seud- 1/sult-1* during the critical window appears to be a conserved molecular decision across environments, natural isolates and sister species. We note however that the other switch gene (*eud-1*) was elevated in Eu-biased strains, and promoters and enhancers of both genes are likely targets for selection.

### Differential expression analysis across diverse conditions reveals shared pathways

After having examined the developmental regulation of genes known to affect mouth-form (*i.e.*, the GRN for mouth-form plasticity), we leveraged our data set to identify additional genes involved in mouth-form plasticity. We combined all previously analyzed samples (PS312 in agar & liquid culture, seven mouth-form mutant strains, four *P. pacificus* strains, and *P. exspectatus* samples) (**Supplementary** Fig. 1A) to build a comprehensive transcriptomic dataset that should filter out all but a core set of mouth-form DEGs. We used k-medoid clustering to separate samples into four developmental clusters (**Fig. 5A**, **Supplementary** Fig. 16) based solely on gene expression (agnostic to sampling time point) to accommodate any discrepancy in developmental rate between different conditions. Cluster 1 contained samples from 0 and 12 h. Cluster 2 contained mostly 24h samples, as well as a few samples from 12h and 48 h. Cluster 3 was almost entirely 48 h samples, and Cluster 4 contained exclusively 60 h and 72 h samples (**Fig. 5A**).

**Figure 5.**
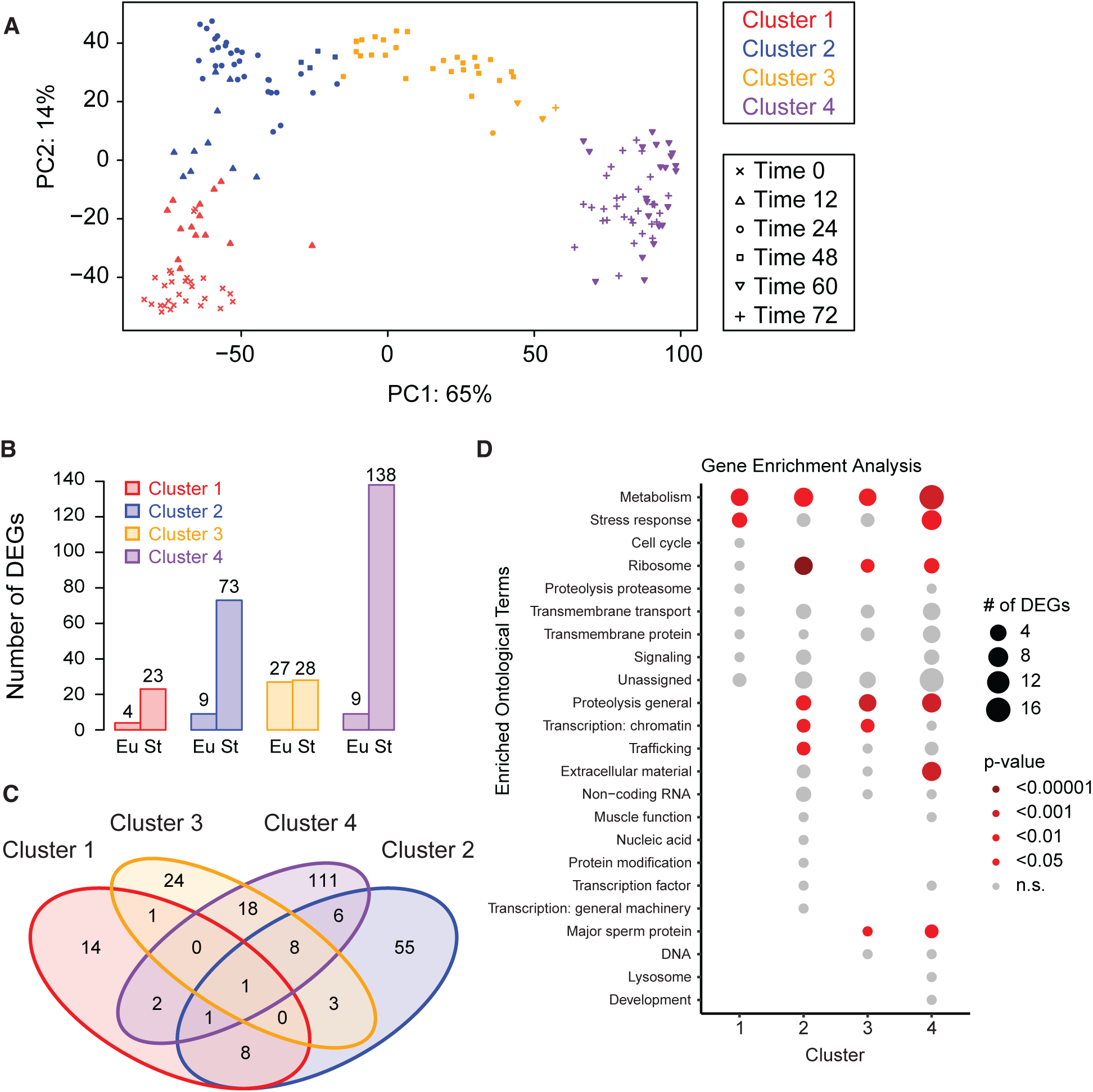
Differential expression analysis across diverse conditions reveals shared pathways. A) PCA and k-medoid clustering of all samples based on gene expression. Color indicates cluster. Shape inidicates timepoint. B) The number of differentially expressed genes (DEGs) between Eu and St samples within each cluster (adj p <0.05, FC >1.5). C) Venn diagram showing the overlap in differentially expressed genes between each cluster. D) Gene enrichment analysis of DEGs within each cluster using WormCat. The size of the bubble corresponds to the number of DEGs in that category and the color corresponds to the p-value.

We then performed a differential expression analysis on all samples, with specific comparisons made between St and Eu samples within each cluster. In general, we found that more genes were upregulated in St worms (*i.e.* down-regulated in Eu worms). Cluster 1 had only 27 DEGs (**Fig. 5B**). Cluster 2 had 82 DEGs, 90% of which were upregulated in St worms. Only Cluster 3 had a roughly equal number of genes that were upregulated in Eu and St worms with a total number of 55 DEGs. Cluster 3 contained almost entirely 48 h samples and represents the middle of the critical window for mouth-form development. This reduction in the number of DEGs at 48 h relative to both earlier and later time points is consistent with our observations in both the environmental comparison (**Fig. 2C**), strain comparison (**Fig. 3C**), and to a lesser extent in the species comparison (**Fig. 5D**). Also consistent with the environmental comparison, we found the largest transcriptomic differences during late larval development and early adulthood, with Cluster 4 having 147 DEGs. Again, most of these (138 genes) were upregulated in St worms. Clusters 3 and 4 shared the greatest number of DEGs; however, the majority of DEGs were unique to a single cluster (**Fig. 6C**). Only one DEG was found in all four clusters (PPA36778) and is most likely a ribosomal protein. Importantly, only two mouth-form genes were significantly differentially expressed in this analysis. The sulfatase *eud-1* showed significant differences in expression between Eu and St worms in Clusters 1 and 3, whereas the sulfotransferase *seud-1/sult-1* showed significant differences in Cluster 3.

**Figure 6.**
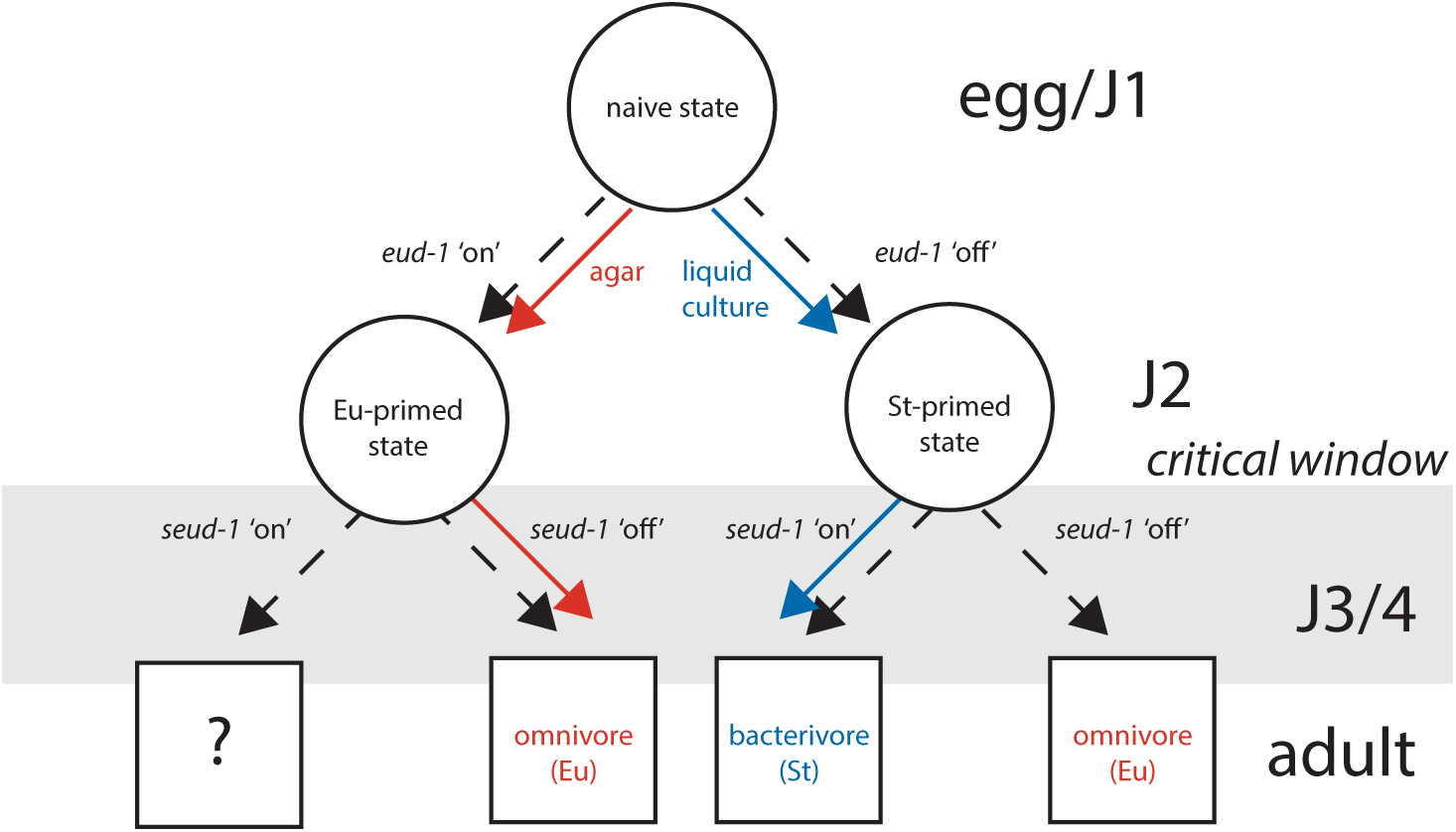
Conceptual model for how switch genes act sequentially across development to regulate mouth form plasticity. Mouth form outcomes given in boxes are based on phenotypes of mutants except for the outcome when both *eud-1* and *seud-1/sult-1* are ‘on’, as that result is unknown.

Thus, even when comparing across multiple inputs to mouth-form development, which effectively filters the total number of DEGs, sulfation comes out as the critical axis of plasticity.

We performed a Gene Enrichment Analysis using WormCat 2.0 (Holdorf et al. 2020) on the *C. elegans* homologs of the DEGs from each cluster to find common pathways affecting morph development (**Supplemental Table 3**). We found significant enrichment of metabolic genes in every cluster (**Fig. 5D**). These metabolic genes included both switch genes (*eud-1* and *seud-1/sult-1*) in Cluster 3 and paralogs of each switch gene in Cluster 4. Within Cluster 4, there was a specific enrichment of genes involved in lipid metabolism. This is consistent with prior comparisons of gene expression between different mouth-form GRN mutants which implicated genes involved in lipid metabolism (Bui and Ragsdale 2019) and starvation response (Casasa et al. 2021) and with the recently identified role of a Mediator homolog in regulating mouth form (Casasa et al. 2023). We also found an enrichment of genes pertaining to extracellular material in Cluster 4, which includes the samples in which the adult mouth is being formed. The tooth-like denticles of *P. pacificus* are cuticular structures composed of chitin (Sun et al. 2022b), therefore the differential expression of collagens and matrix-secreting genes between Eu and St worms suggests that distinct extracellular matrix components contribute to the formation of each mouth form.

This compositional difference could be due either to the presence/absence of certain components or their relative quantities. In summary, comparing gene expression across multiple conditions highlighted metabolism, stress response, ribosomal function and chromatin regulation as shared processes which may be involved in regulating mouth-form development.

## DISCUSSION

The environmental triggers and resulting phenotypes for several model systems of plasticity have been described over the last two decades, contributing to the appreciation of plasticity in development as a potential source of evolutionary novelty (Moczek et al. 2011; Simpson et al. 2011; Levis et al. 2018; Sommer 2020). However, the molecular mechanisms that mediate plasticity remain unknown beyond a few isolated genetic factors, limiting the incorporation of plasticity into standard evolutionary theory (Laland et al. 2015). Recently, genetic studies in *P. pacificus* have laid the groundwork for a gene regulatory network for mouth-form development. Here, we describe the temporal regulation of the mouth-form genes in the context of both environmental induction and different genetic backgrounds. When interpreted in the context of the critical window (Werner et al. 2023) these data allow us to evaluate the regulatory logic of plasticity.

Our results highlight four tenets of genetic regulation of mouth-form plasticity. First, we found that only a subset of the GRN is differentially expressed by environmental conditions. This indicates that not all genes which have dominant effects on a plastic trait are involved in its environmental induction.

However, it is formally possible that other conditions could reveal different patterns of environmental induction. Moreover, only two mouth-form switch genes were expressed during the critical window for mouth-form development. This surprising finding argues that of the many genes that compose a GRN, there may be only 1-2 major switch genes controlling the developmental decision of a plastic trait.

Second, transcriptional regulation of the GRN is largely limited to discrete time points. When upstream genes are mutated or overexpressed, the effect on downstream genes is (with some exceptions) confined to their wild-type time points of regulation, indicating that nodes in plasticity networks are under temporal regulatory constraint.

Third, regulatory circuits exist between enzymes with different functions (*e.g. nag-1/2*, *eud-1*, *seud- 1/sult-1*, *lsy-12*, and *mbd-2*). Although the enzymatic roles for each of these genes are not yet known in *Pristionchus*, it seems likely that this circuit is connected by signaling molecules which are either directly or indirectly linked with the abundance of their substrates and products. *eud-1* and *seud-1/sult-1* work by removing or adding sulfate groups to some unknown substrate (Igreja and Sommer 2022). Sulfation is an important biochemical mechanism for regulating the activity of signaling molecules, metabolizing xenobiotics, and generating diverse extracellular ligands (Strott 2002). Our results underscore its importance as a key switch mechanism for mouth-form plasticity.

Finally, fourth, the two major switch genes appear to act sequentially. The observed pattern of differential expression leads to a model in which elevated expression of *eud-1* prior to and during the critical window sets an initial trajectory for mouth-form development. *seud-1/sult-1*’s expression in J3 and J4 larvae provides a second check point closer to the close of the critical window, with binary mouth form outcomes depending upon the balance between *eud-1* and *seud-1/sult-1*’s expression. This conceptual model is reminiscent of a binary decision diagram (Akers 1978), ultimately yielding a Boolean outcome: omnivore (Eurystomatous) or bacterivore (Stenostomatous)(**Fig. 6**).

It will be interesting to see if these GRN patterns can be found in other systems of plasticity. Beyond the known GRN, we also found that metabolic genes were differentially expressed between Eu and St worms throughout development, corroborating previous results that mouth-form is linked with metabolism in *Pristionchus* (Bui and Ragsdale 2019; Casasa et al. 2021). Notably, this induction begins prior to the critical window, and may therefore feed into the regulation of switch gene transcription. Diet and metabolic state are known regulators of developmental plasticity in many established systems of phenotypic plasticity (Pfennig 1990; Emlen 1994; Moczek 1998; Simpson et al. 2011), and yet the molecular mechanisms of the metabolic regulation of mouth-form plasticity remains unknown.

Altogether, our results reveal the temporal and regulatory logic of mouth-form plasticity in *Pristionchus* and provide a testable set of hypotheses for other plastic traits and species.

## METHODS

### Nematode strains and maintenance

The reference *P. pacificus* strain PS312 (RRID:WB-STRAIN:WBStrain00047433) was used for the environmental samples (agar plates and liquid culture). All other strains were grown only on agar plates. The following mutants were used: *Ppa-eud-1*(*tu1069*)*, Ppa-Ex[eud-1]*, ( *Ppa-lys-12* (*tu319*), *Ppa-sult-1* (*tu1061*), *Ppa-nag1/2* (*RS3195/tu1142/tu1143*), *Ppa-nhr40* (*tu505*), and *Ppa-mbd-2* (*tu365*). The *eud-1* mutant is a novel CRISPR/Cas9 induced 10 bp deletion in exon 3, while all other mutants were previously published ethyl methanesulfonate (EMS) or CRISPR/Cas9 induced mutations (**Table 1**).

However, an inspection of the sequencing reads from the *sult-1* mutant line revealed that despite having a mutant phenotype, this line does not contain the reported frameshift mutation: it has a 9 bp deletion rather than the previously reported 10 bp deletion in the 9^th^ exon (**Supplemental Fig. 17,** Namdeo *et al*. 2018). Four strains of *P. pacificus* (RS5410, RS5427, RSC017, and RSA100) and a strain of *P. exspectatus* (RS5522B) were used to assess the role of genetic background on the mouth-form GRN. For general maintenance of nematode cultures, five young adults were passed every 4–6 days on 60 mm NGM-agar plates at 20 °C seeded with 300 µl of overnight cultures of *Escherichia coli* OP50 (grown in LB medium at 37 °C) and covered with parafilm. Majority hermaphrodite (>95%) cultures were used for all experiments.

### Phenotyping of *daf-12* mutants

We phenotyped worms from two different mutant lines of the nuclear hormone receptor *daf-12* (**Supplemental Fig. 18**). RS2209 (*tu381*) has a mutation in the splice acceptor of the hinge region and RS2272 (*tu390*) has a premature stop codon in the ligand binding domain (Ogawa et al. 2009). We grew worms on NGM agar plates and determined the mouth-form ratio (% Eu) of each strain and wildtype PS312. Error bars represent S.E.M. for n=3 independent biological replicates, with >20 worms phenotyped per replicate.

### RNA-Sequencing

Developmental RNA-seq was performed on worms from different strains and grown in different environmental conditions (liquid culture and NGM agar plates), at the time-points described in **Fig. 1C & Supplemental Fig. 1A**. Details regarding the culture state, sampled time points, number of replicates and strain IDs for each condition are given in **Supplementary Table 1**. To synchronize worms we performed cuticular disruption of gravid adults with bleach/NaOH (Stiernagle 2006) and aliquoted eggs-J1s to NGM-agar plates or S-medium liquid culture OP50 *E. coli* (Werner et al. 2017), or for t’=0, directly resuspended in 500 µl Trizole (Ambion cat. #15596026). Thus, the agar t’=0 timepoint represents the start of both the agar and the liquid culture experiments for the environmental comparison. Liquid cultures were incubated at 180 rpm, 20-22 °C. Worms at different stages were then collected at corresponding time points by washing from plates with M9, or simply decanting from liquid culture, and filtering through 5 µM filters. Washed worms were then resuspended in 500 µl Trizole and freeze-thawed 3x between 37 °C and a small liquid nitrogen container to disrupt the cuticle. RNA was then extracted following the manufacturer’s protocol and purified using Zymo RNA clean & concentrator-25 columns (Zymo, cat. # R1017). RNA was then eluted in 50 µl water and quantified by nanodrop (A260/280). 500 ng to 1 µg of total RNA was converted to Illumina sequencing libraries using the NEB Ultra II Directional RNA Library Prep kit with Sample Purification Beads (NEB cat. #E7765), with 10-14 PCR cycles (determined empirically to achieve the minimum number of cycles per library). Libraries were then sequenced on an Illumina HiSeq 2000 in single-end mode, except for the 12-hour time points, which were sequenced in paired-end mode. We kept samples with a minimum of 3 million mapped reads and average quality scores greater than 30.

### Read Mapping

We used Salmon 1.3.0 (Patro et al. 2017) to map *P. pacificus* sequencing reads to the *P. pacificus* transcriptome (El Paco annotation, version 2) and *P. exspectatus* reads to the reference transcriptome (Rödelsperger et al. 2014). We first indexed the El Paco transcriptome, and then quantified the sequencing reads against this index, correcting for GC bias.

### Differential Expression Analysis

We imported the quantified reads to Rstudio using tximport and performed differential gene expression analysis using DESeq2 (Love et al. 2014). We first did a differential expression analysis on each condition (agar, liquid culture, individual strains, individual mutant lines, *P. exspectatus* samples) to check that the replicates from the same time points clustered together (**Supplemental Fig. 2-7**). We then moved forward with the analysis and compared gene expression across different conditions.

We performed separate analyses for each of the different conditions and compared samples at the same time point using DESeq2 (design= ∼MF_time) for environmental and genetic background comparisons and DESeq2 (design= ∼condition) for mutant comparisons. We did not impose a fold change threshold but considered all genes with an *adjusted p-value*<0.05 to be significantly differentially expressed.

For the species comparison, we identified genes that were 1:1 orthologs through reciprocal protein blasting with an e-value of 1e-3. This resulted in 11,556 1:1 orthologs between the two species. We found more orthologs when querying *P. exspectatus* against *P. pacificus*, but the number was greatly reduced when we queried *P. pacificus* proteins against *P. exspectatus*. However, this pipeline excluded both *mbd- 2* and *eud-1*, two genes which should be 1:1 orthologs. The problem appeared to be the poor *P. exspectatus* annotation. We were able to identify the *P. exspectatus* annotation for *eud-1* and add it back into the dataset, but we could not identify the *P. exspectatus* annotation for *mbd-2*, so it was left out of the species comparison. We then gave the *P. exspectatus* reads the *P. pacificus* ortholog’s gene ID and performed a differential expression analysis between species by timepoint using DESeq2 (design = ∼ MF_time). This resulted in a comparison of 11,504 transcripts with a nonzero total read count, compared to 22,631 transcripts in the initial species comparison.

To identify a core list of genes that are differentially regulated in St and Eu worms, we performed a differential expression analysis on all samples from the different conditions together. We first ran DESeq2 on all samples with replicates grouped such that the comparison was between all unique sample types (*i.e.* agar samples from the 0 time-point were compared against all other samples, including other agar samples). We performed a principal component analysis on the log2-transformed results and noted that the first principal component explained 65% of the variance in gene expression (**Figure 6A**). The samples separated along this principal component according to time point. We plotted the dissimilarity between samples against the number of clusters and noted diminishing returns when dividing the samples between more than four clusters (**Supplemental Fig. 18**). We then performed k-medoid clustering to subset the samples into four clusters. Finally, we ran DESeq2 on all samples, with comparisons being made between Eu and St samples within each cluster (design=∼MF_clust). This produced a list of differentially expressed genes for each cluster with fold change threshold of 1.5 (lfcthreshold = 0.585) and >95% confidence (*adjusted p-value* <0.05).

### Gene Enrichment Analysis

We used blastp to identify *C. elegans* orthologs and WormBase IDs of the *P. pacificus* genes that were differentially expressed between Eu and St worms. Only 53% (134 genes) of the *P. pacificus* DEGs from the combined analysis had a blast hit. Gene enrichment analysis was completed using WormCat 2.0 (Holdorf et al. 2020) on the WormBase IDs.

### Shiny App

We made a shiny app in R (version 4.3.3) that plots the normalized expression of any *Pristionchus* gene in our different conditions across development. The app extracts the values from a spreadsheet which corresponds to the All_samples_ntd tab in **Supplemental Table 4**. The custom code used to generate the app can be found at: https://github.com/drjuliejung/GRN-shiny.

### Data access

The RNA sequencing data generated in this study have been submitted to the NCBI BioProject database (https://www.ncbi.nlm.nih.gov/bioproject/) under accession number PRJNA628502.

## Acknowledgments

We thank Thomas King and Audrey Brown for thoughtful critique of the manuscript. This study was funded by the Max Planck Society and NIH grant R35GM150720-01 to M.S.W.

## Author contributions

M.S.W. and R.J.S. conceived and designed experiments for sampling and sequencing RNA. M.S.W conducted RNAseq with assistance from T.L. S.R. performed all bioinformatic analyses with assistance from M.S.W. J.J. produced the Shiny app from the data. S.N. phenotyped *daf-12* mutants. S.R. wrote the manuscript with assistance from M.S.W.

## Notes

https://pristionchus-transcriptomics.shinyapps.io/GRN-shiny/

### Competing Interest Statement

The authors have declared no competing interest.

