## Supplemental Figures for "Developmental transcriptomics in *Pristionchus* reveals the logic of a plasticity gene regulatory network"

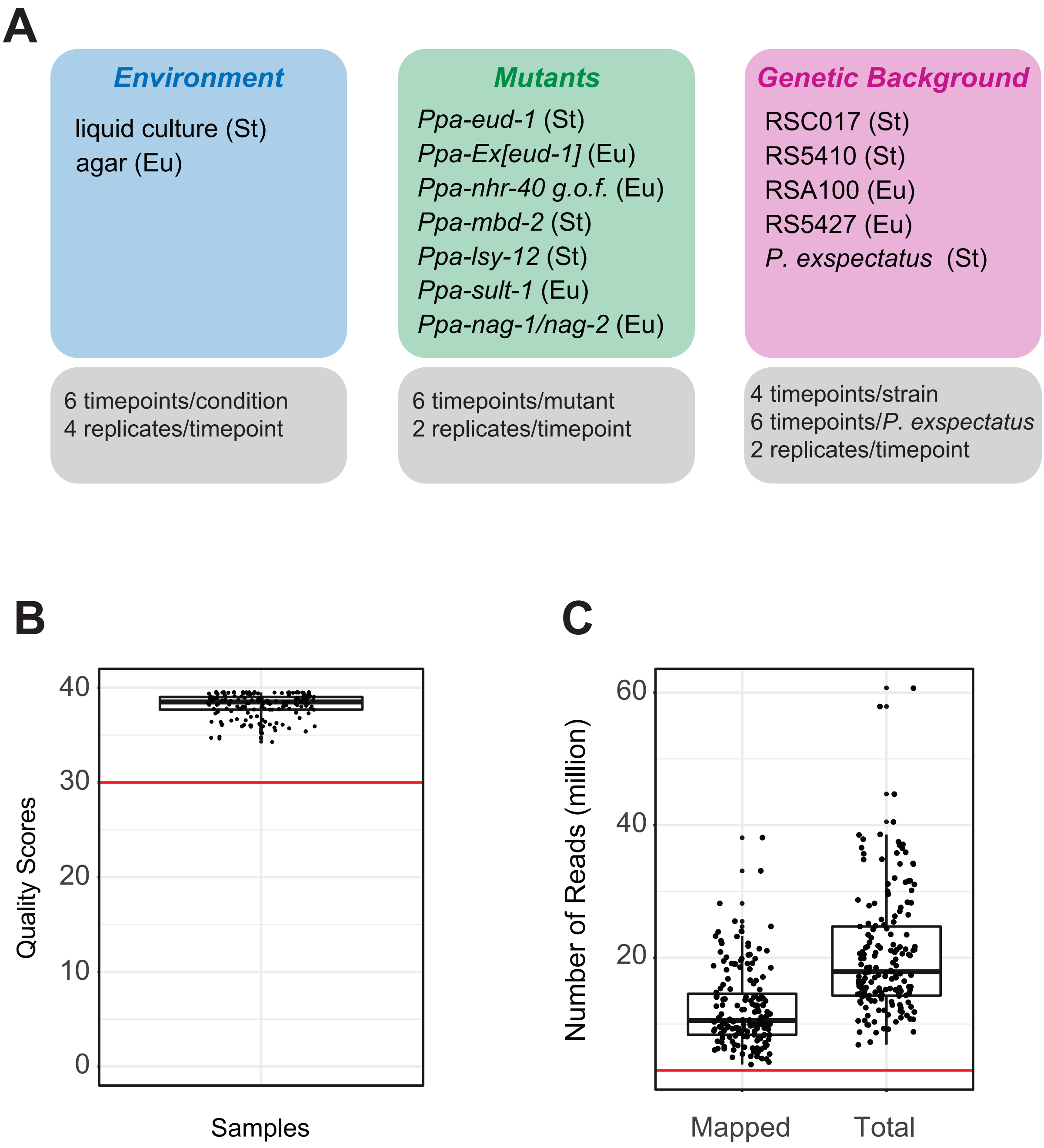


**Supplemental Figure. 1. Experimental design and RNAseq quality scores**
A) Table of conditions and worm lines sampled for RNAseq along with timepoints and number of replicates.
B) Quality scores of RNAseq data. Red line represents threshold of 30.
C)Total number of RNAseq reads and number of reads that mapped to transcriptome. Red line represents threshold for sample inclusion of 3 million reads.

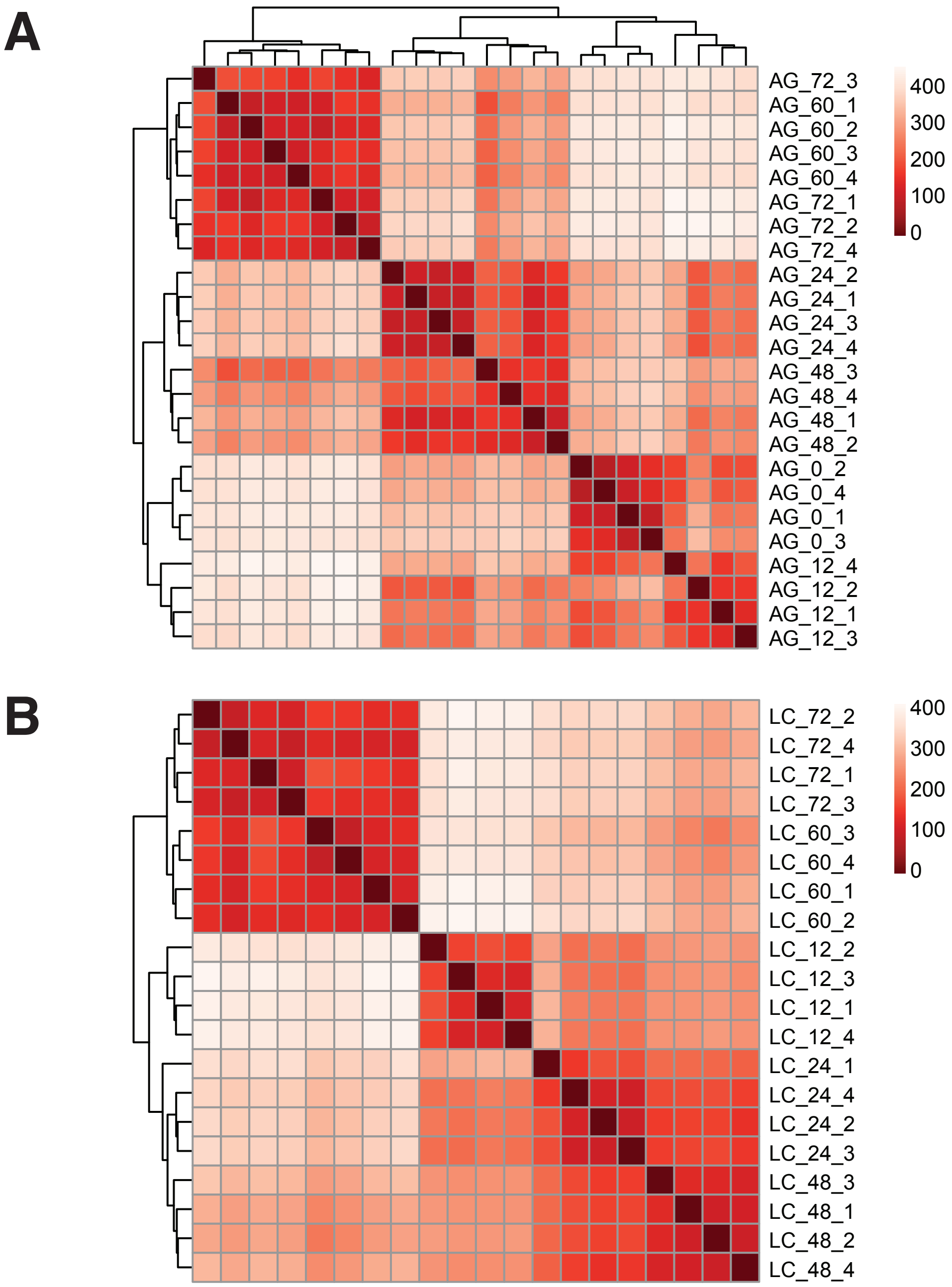


**Supplemental Fig. 2.** Heatmaps of normalized gene expression from A) six developmental timpoints grown on agar plates and B) five developmental timepoints grown in liquid culture. The zero timepoint represents eggs which were aliquoted between agar and liquid cultures.

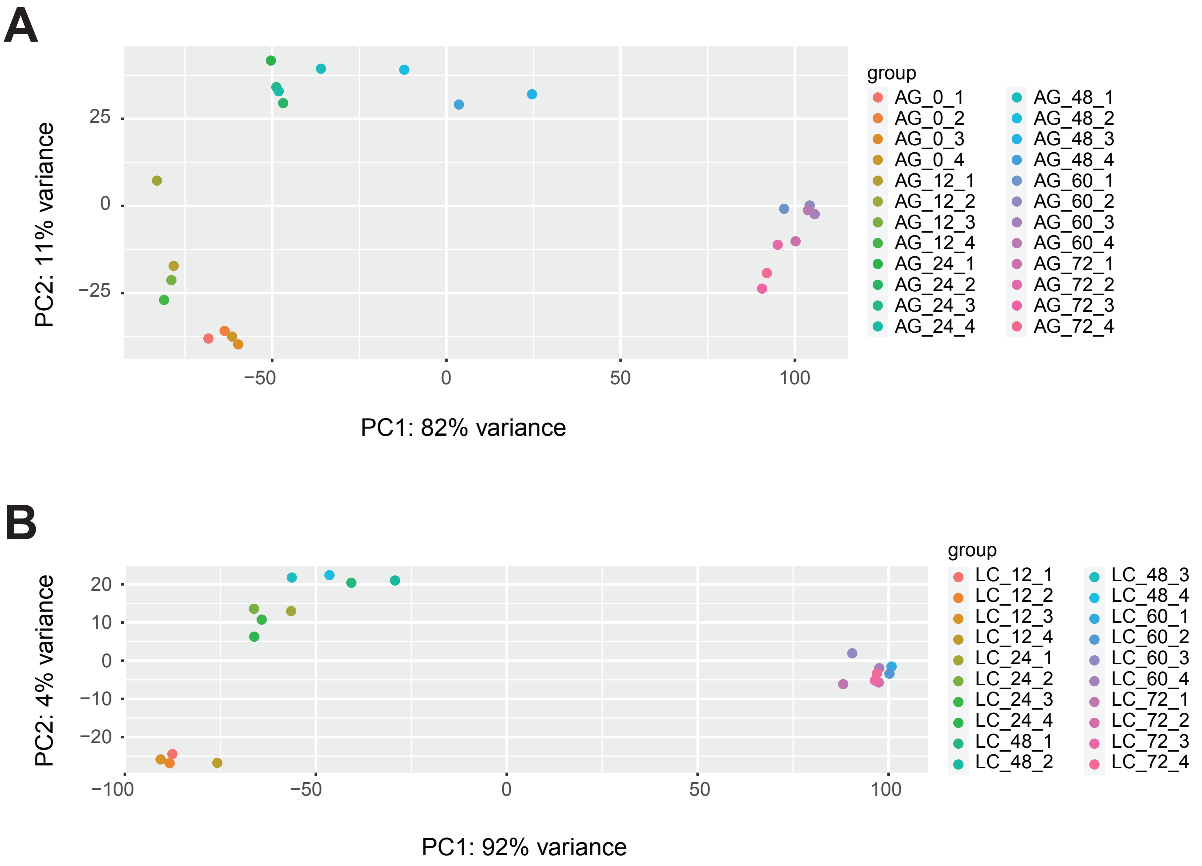


**Supplementary Fig. 3.** Principal Component Analysis (PCA) of normalized gene expression from A) eggs and worms grown on agar plates and B) worms grown in in liquid culture.

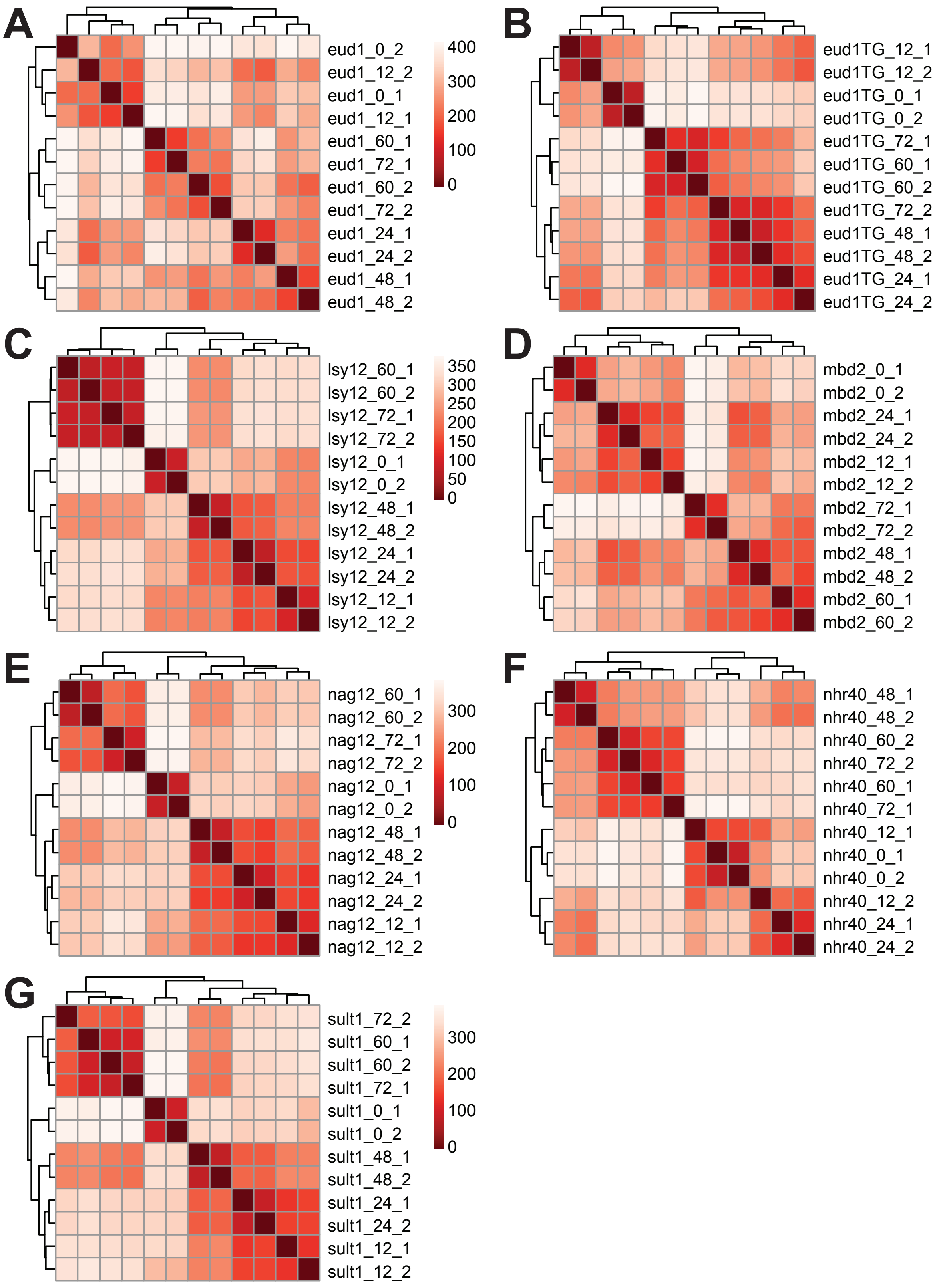


**Supplementary Fig. 4.** Heatmaps of gene expression from eggs and five developmental time points from seven mutant strains: A) *Ppa-eud-1* B) *Ppa-Ex[eud-1]* C) *Ppa-lsy12* d) *Ppa-mbd2* E) *Ppa-nag1/nag2* F) *Ppa-nhr40 gof* and G) *Ppa-sult-1*.

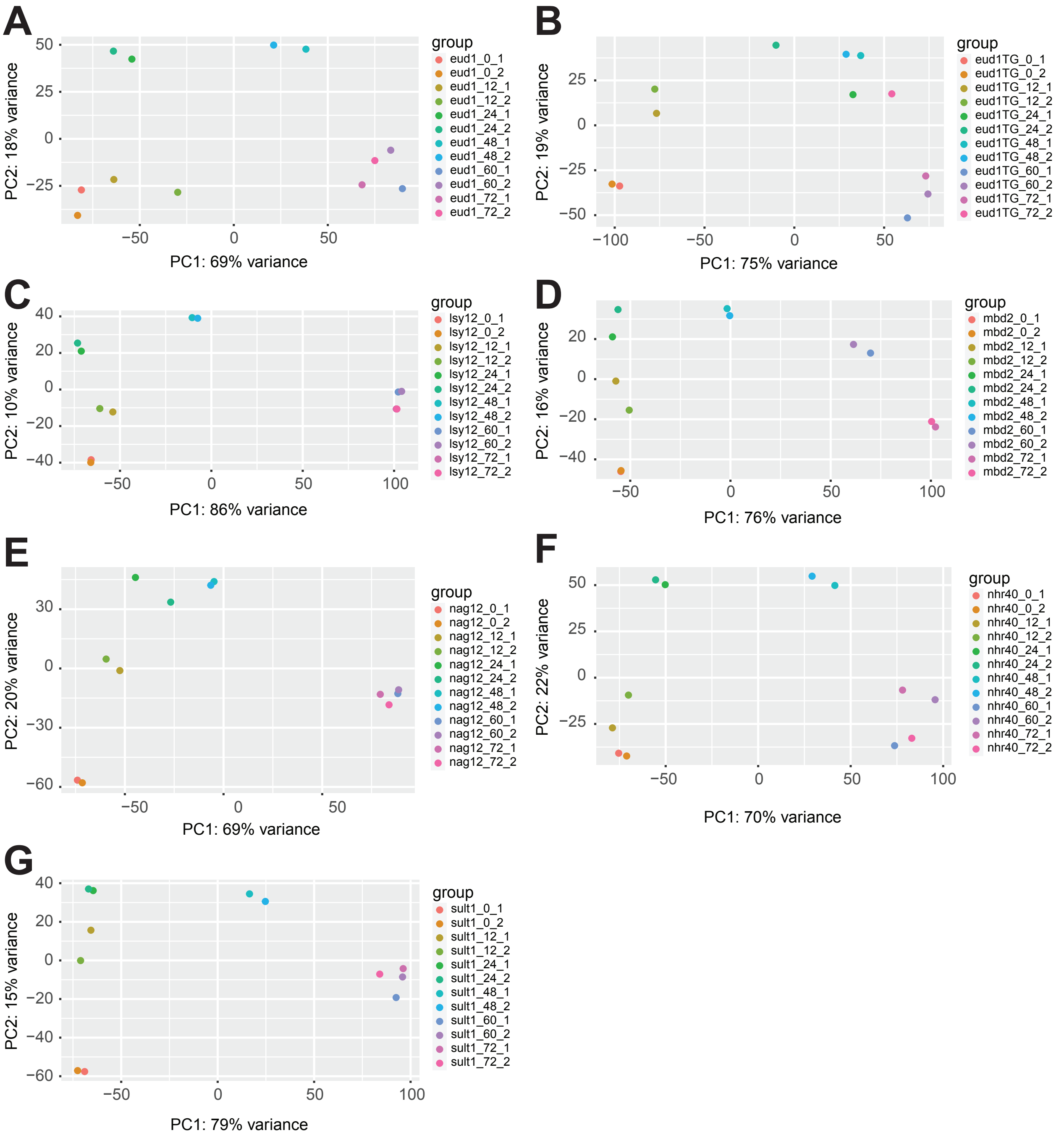


**Supplementary Fig. 5.** Principal Component Analysis (PCA) of normalized gene expression from seven mutant strains: A) *Ppa-eud-1* B) *Ppa-Ex[eud-1]* C) *Ppa-lsy12* D) *Ppa-mbd2* E) *Ppa-nag1/nag2* F) *Ppa-nhr40 gof* and G) *Ppa-sult-1.*

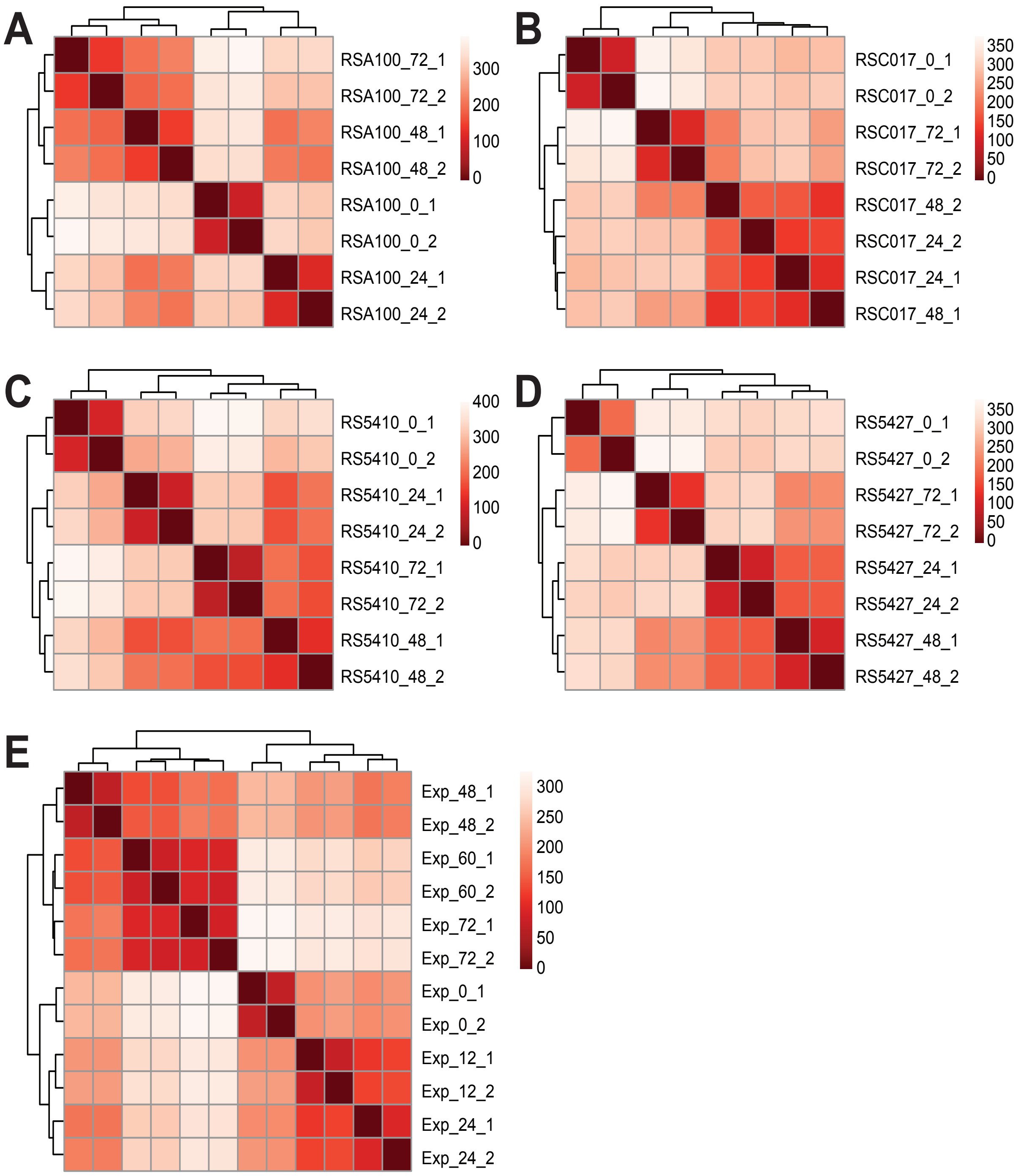


**Supplementary Fig. 6.** Heatmaps of normalized gene expression from four *P. pacificus strains* A) RSA100, B) RSC017, C) RS5410, D) RS5427 and from E) *P. exspectatus*.

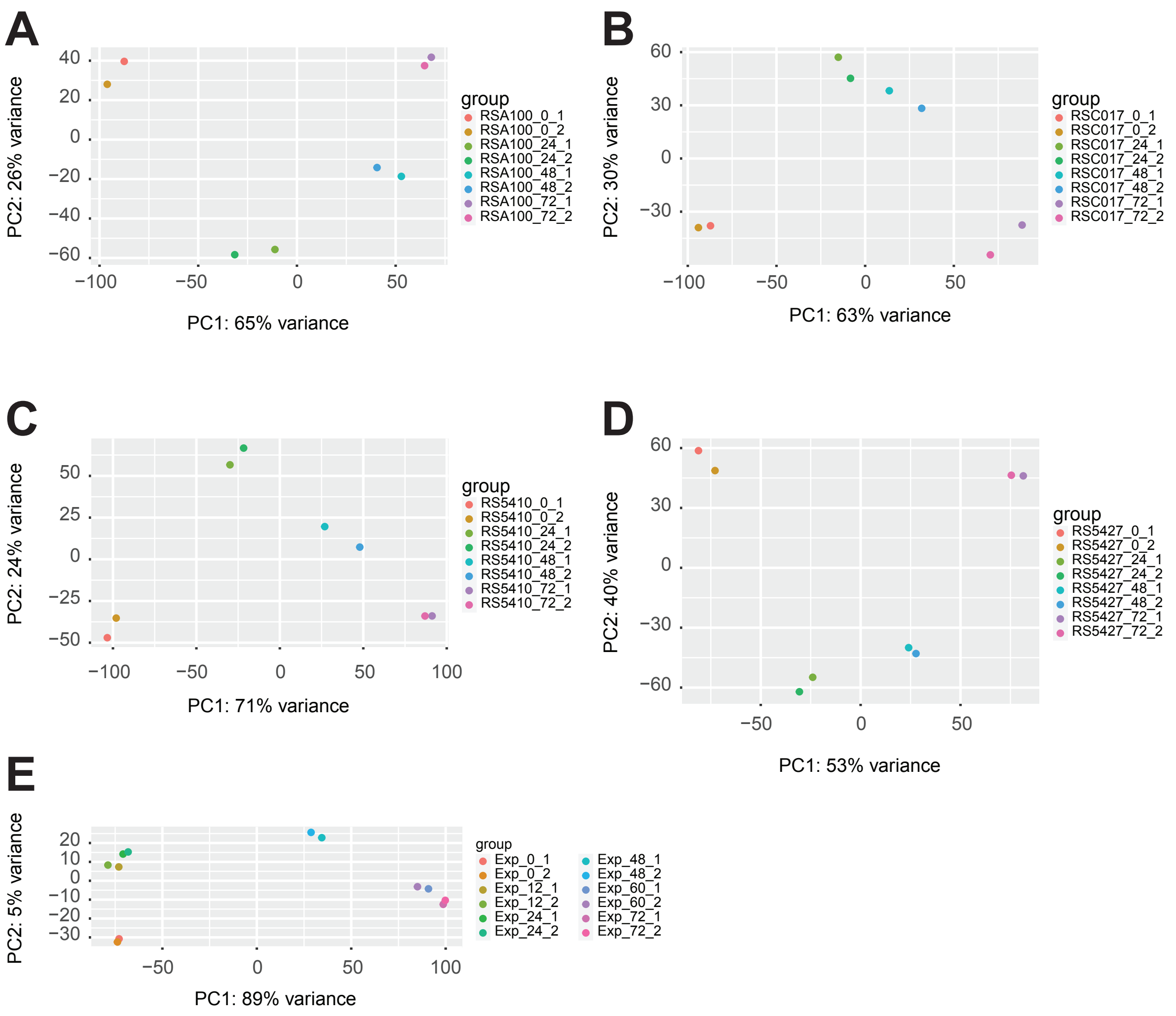


**Supplementary Fig. 7.** Principal Component Analysis (PCA) of normalized gene expression from from four *P. pacificus strains* A) RSA100, B) RSC017, C) RS5410, D) RS5427 and from E) *P. exspectatus*.

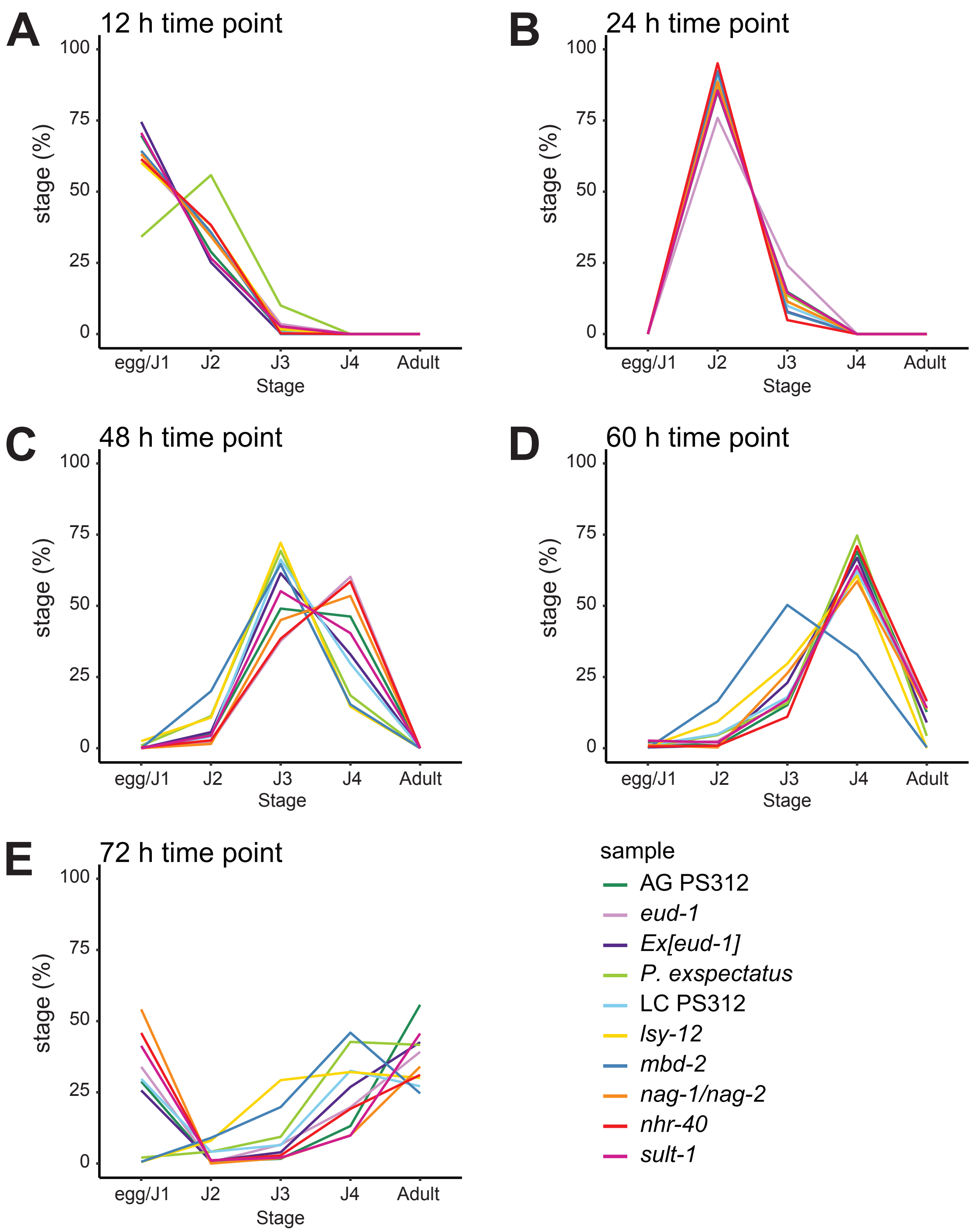


**Supplemental Fig. 8.** Developmental stage of worms at from environmental, mutant, and *P. exspectatus* cultures at A) 12 h B) 24 h C) 48 h D) 60 h E) 72 h time points.

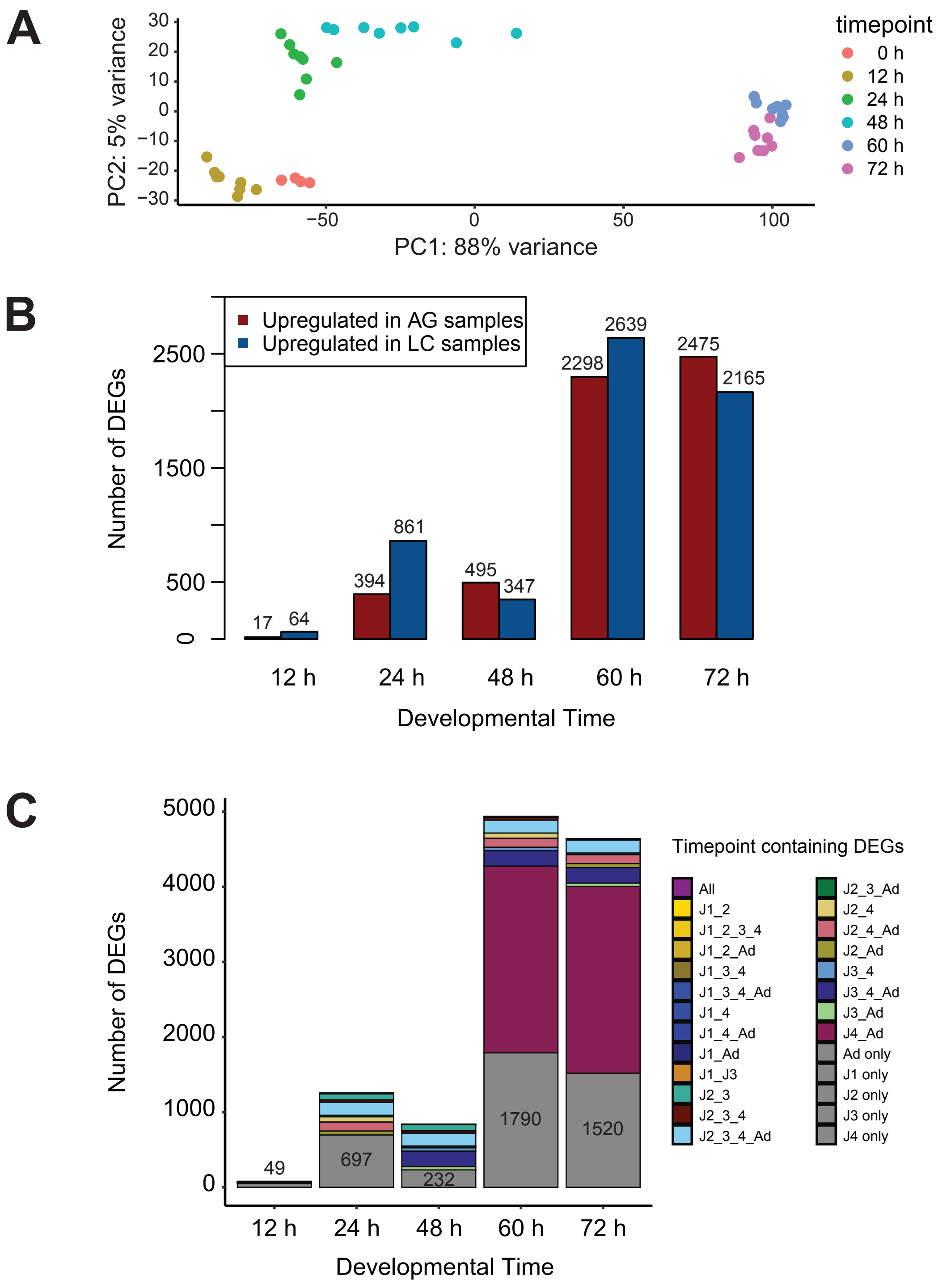


**Supplemental Fig. 9.** Additional analyses from the environmental comparison. A) Principal Component Analysis of normalized gene expression results from PS312 grown on NGM agar plates and S-media liquid cutlures with samples colored by time point. B) Number of differentially expressed genes (DEGs) at each sampled time point upregulated in agar (AG, red) or liquid culture (LC, blue). C) Number of differentially expressed genes (DEGs) at each sampled time point with DEGs shared between time points colored the same. The exception is the gray blocks, which represent the number of DEGs that were unique to each time point.

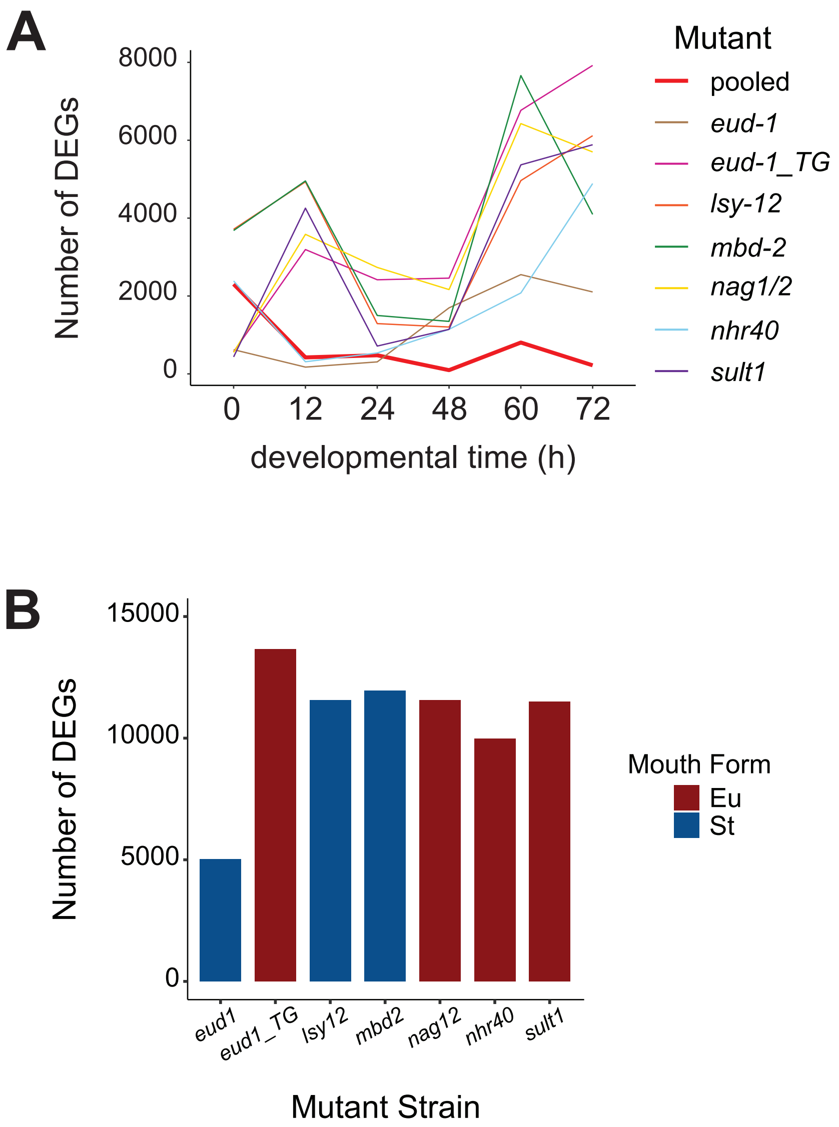


**Supplemental Fig. 10.** Transcriptomic comparisons between mouth-form mutants and PS312 grown on agar plates. A) Number of differentially expressed genes (DEGs) per time point between each mutant strain and PS312. Red line shows the number of DEGs between pooled Eu and St mutants. B)Total number of unique differentially expressed genes across all time points for each mutant  line comapred to wildtype PS312 (*adjusted p-value* <0.05). Bars are colored by mouth-form of mutant strain. *eud1_TG* is the overexpression line *Ppa-Ex[eud-1]*.

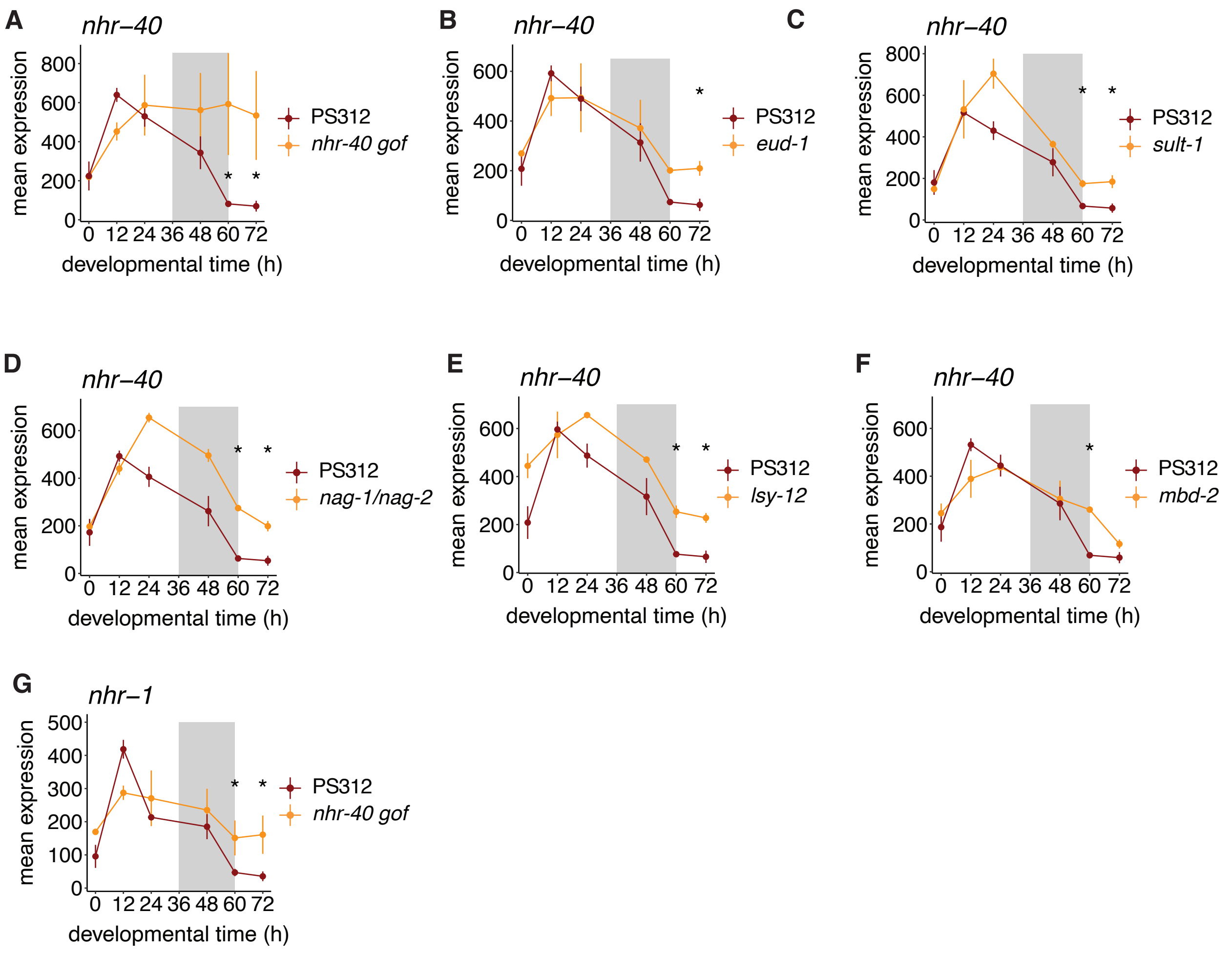


**Supplemental Fig. 11.** Expression of nuclear hormone receptors is affected by mutating mouth-form genes. Plots show mean expression of normalized counts from DESeq2.

**A**

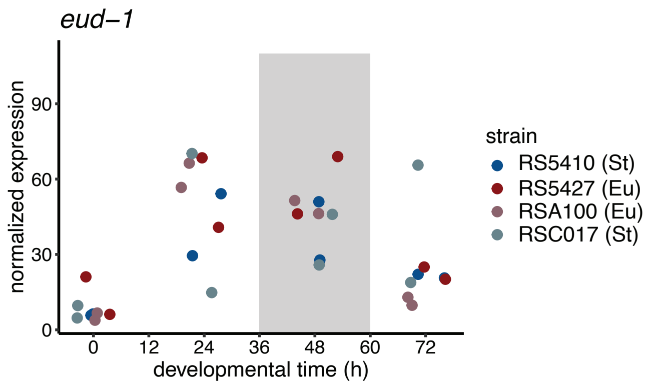


**B**

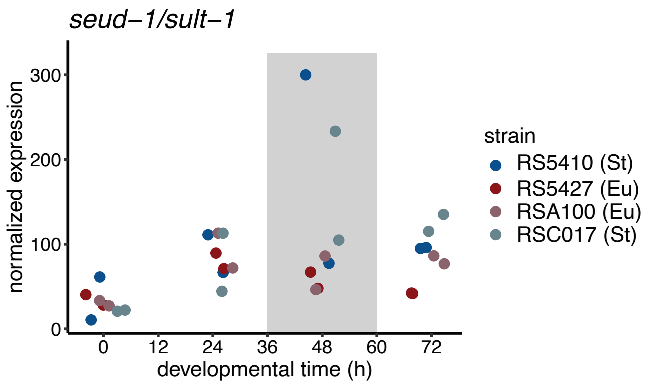


**Supplemental Fig. 12.** Expression of mouth form switch genes (A) *eud-1* and (B) *seud-1/sult-1* in four different natural isolates of *P. pacificus*. Mouth-form bias of strain is given in parentheses.

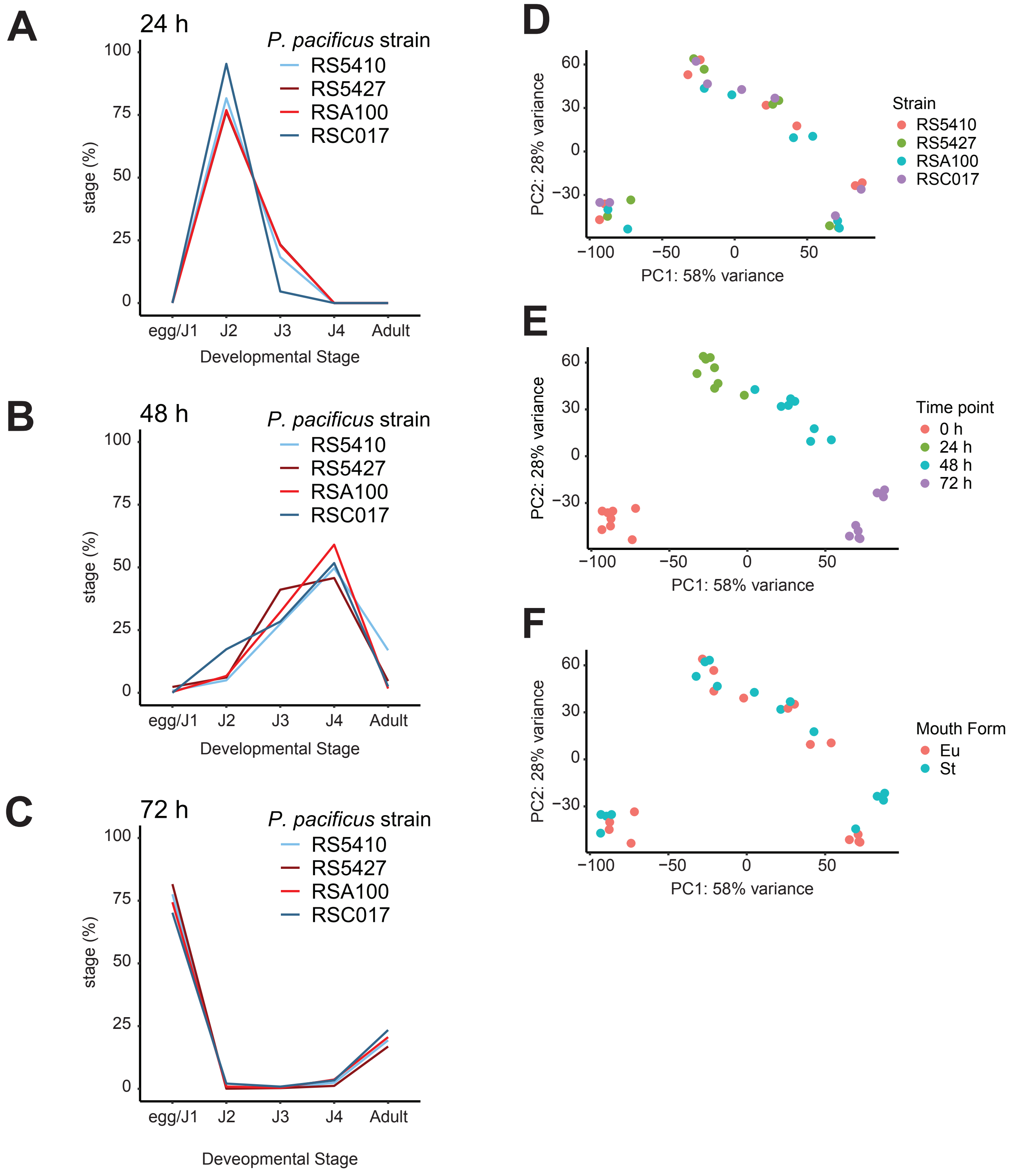


**Supplemental Fig. 13.** Developmental staging of Eu- and St-biased *P. pacificus* strains showing stages present at A) 24 hr, B) 48 hr, and C) 72 hr time points. Principal Component Analysis (PCA) of normalized gene expression colored by D) strain ID, E) sampling time point, and F) mouth-form bias.

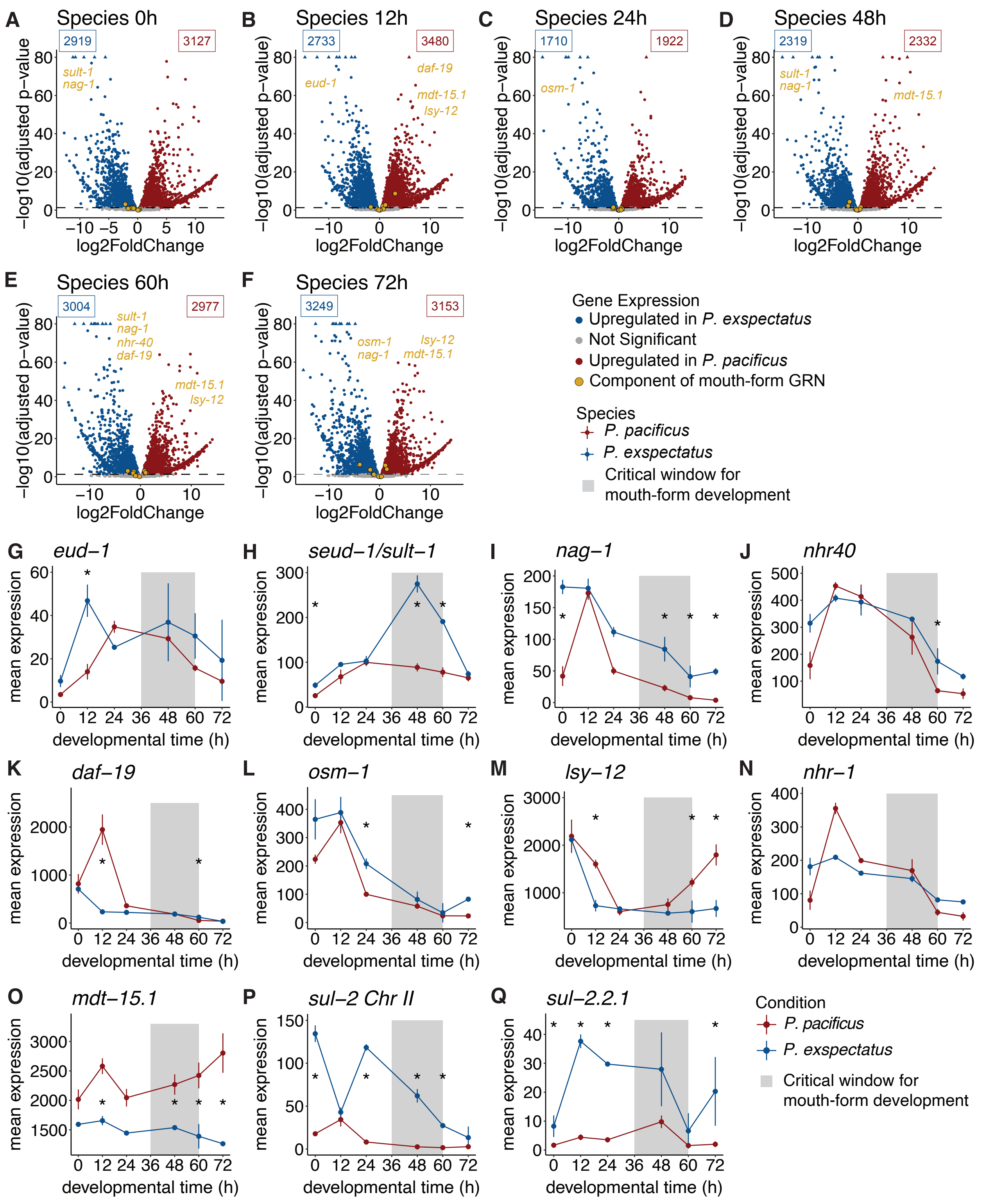


**Supplemental Fig. 14.** *Pristionchus* species exhibit large transcriptomic differences. A-F) Volcano plots of differentially expressed genes (DEGs) between 1:1 orthologs of *P. pacificus* (Eu-biased) and *P. exspectatus* (St-biased) at six developmental time points. Significant DEGs (*adjusted p-values <0.05*) are colored by whether they are upregulated in *P. exspectatus* (blue) or *P. pacificus* (red). Mouth-form genes are colored in gold and labeled on plot when significant. G-Q) Mean expression of normalized counts from DESeq2 for mouth-form genes in two species at six developmental time points. * time points with significant differences in expression (*adjusted p-value* <0.05).

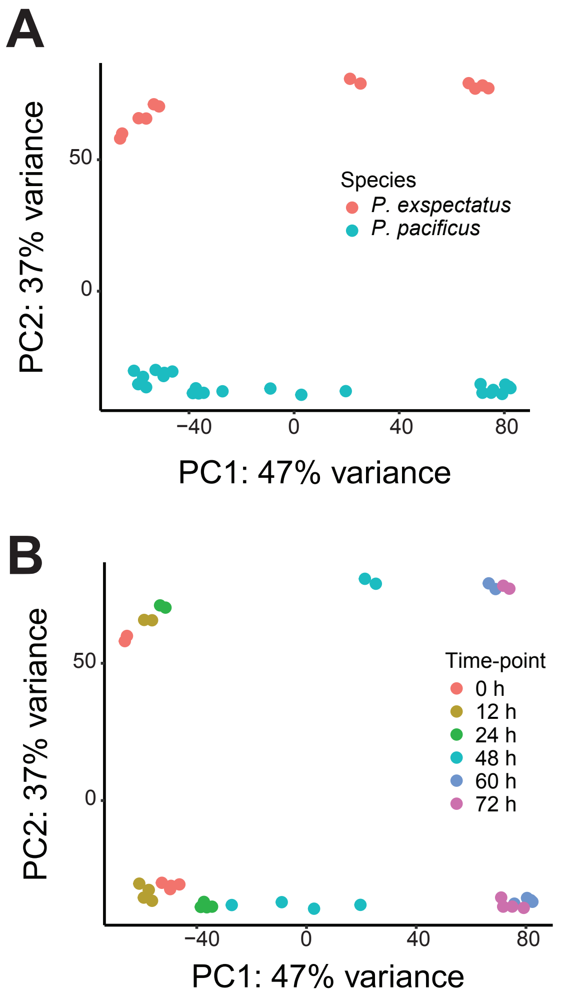


**Supplemental Fig. 15.** Species differences explain substantial amount of variance in gene expression between *P. pacificus* and *P. exspectatus*. Principal component analysis of normalized differential gene expression results from the species comparison using reads from 1:1 orthologs between the species and each species' transcriptome. Each dot represents a different sample, and samples are colored by A) species and B) sampling time point.

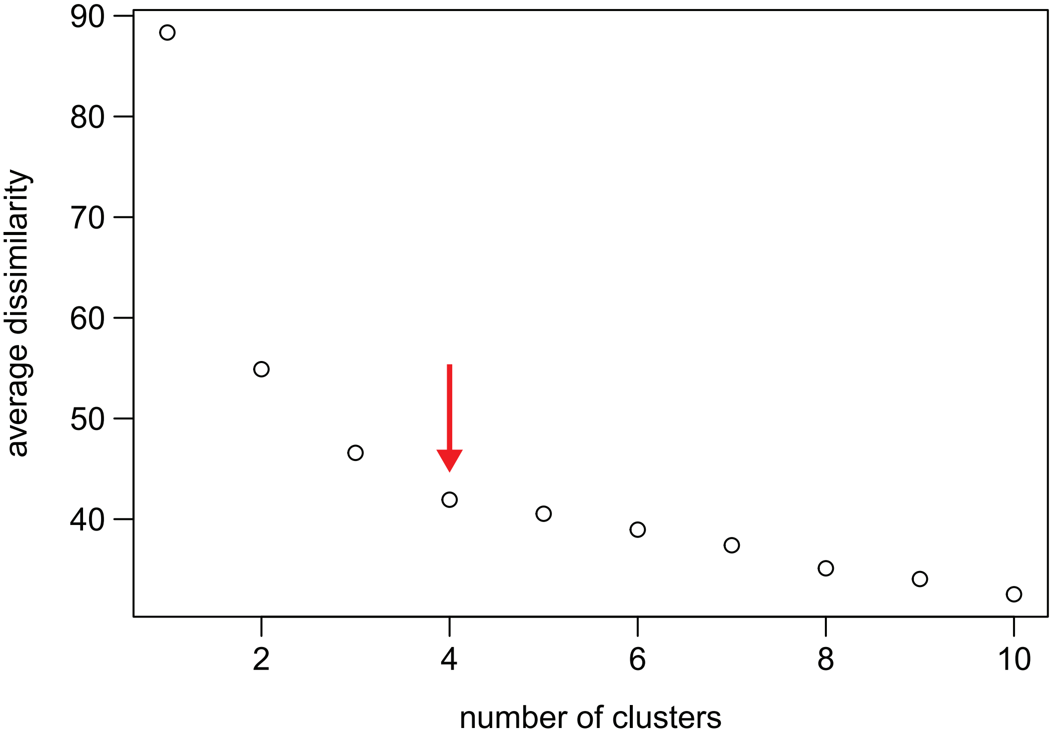


**Supplemental Fig 16.** Dissimilarity analysis used to determine number of clusters. Dissimilarity plot showing the average dissimilarity within a cluster when samples were divided into a given number of clusters. We decided on 4 clusters (indicated by red arrow) based on the declining reduction of dissimilarity when samples were divided into 4 or 5 clusters.

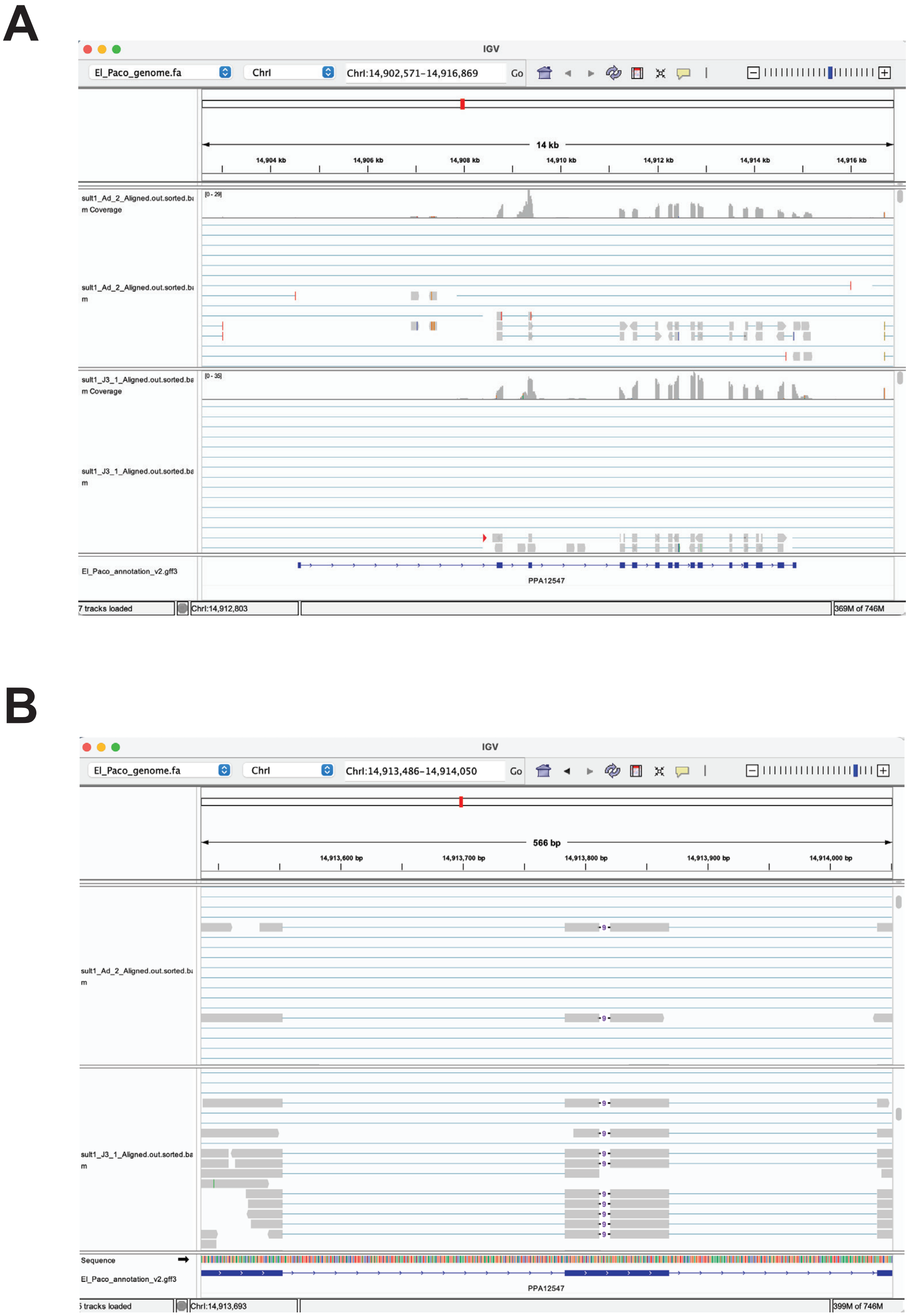


**Supplemental Fig. 17.** Mapping reads from sult-1 samples to genome to check location of the mutation. A) Screenshot of window from IGV showing the entire *seud-1/sult-1* locus and the counts of reads in two different *sult-1* samples aligned across the locus. B) Zoomed in screenshot of window from IGV showing the 9 bp deletion in the 9th exon of *seud-1/sult-1*.

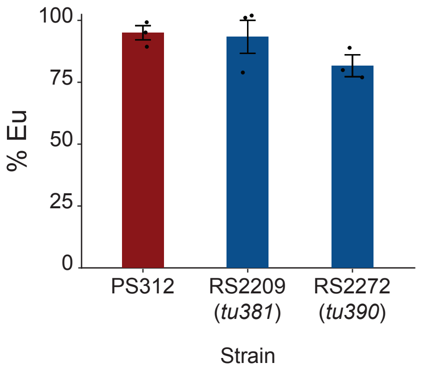


**Supplemental Fig. 18.** Mouth-form phenotype of two different *daf-12* mutants and wild-type PS312 grown on NGM agar plates. Error bars represent S.E.M. for n=3 independent biological replicates, with >20 worms phenotypes per replicate.
